# Comparative morphology of the dormouse skull

**DOI:** 10.1101/2021.04.23.441073

**Authors:** Jesse J. Hennekam

**Affiliations:** Naturalis Biodiversity Center, Leiden, The Netherlands; Hull York Medical School, University of York, United Kingdom

**Keywords:** Dormice, Gliridae, Geometric Morphometrics, Allometry, Skull Morphology, Functional Morphology

## Abstract

Dormice exhibit both inter- and intrageneric variation in cranial and mandibular morphology. Using geometric morphometrics, the form of eight out of nine extant dormouse genera was analysed, to provide a better understanding of the overall morphological variation present within Gliridae. Three genera representing the subfamilies, were studied in more detail. Species-, genus- and family-specific morphological trends are linked with certain habitats and feeding strategies. Smaller dormice show adaptations to a more arboreal lifestyle such as a relatively enlarged braincase and an inferiorly reoriented foramen magnum. Larger dormice show cranial modifications, including clear flattening of the skull and a more posteriorly positioned foramen magnum, hinting towards a more rupicolous lifestyle. Furthermore, specimens inhabiting arid areas appear to have more inflated auditory bullae, whereas other variable features, such as the length of the incisive foramen, were not associated with either size changes nor climatic variables. Lastly, more robust and horizontally orientated zygomatic arches as well as increased robusticity of the molar row appear to be linked with herbivory in dormice, whereas thinner arches and small concave molars are seen in more insectivorous species. This study results in a better understanding of ecological drivers underpinning the morphological divergence present within Gliridae.

## Introduction

Gliridae is considered one of the oldest extant rodent families (Fabre et al., 2012) and consists of nine dormouse genera split over three subfamilies; Glirinae (*Glirulis, Glis*), Leithiinae (*Chaetocauda, Dryomys, Eliomys, Muscardinus, Myomimus* and *Selevinia*), and the monogeneric Graphiurinae (*Graphiurus*) (Holden-Musser et al., 2016). Dispersed widely throughout Africa and Eurasia, the family occupies various ecological habitats, including tropical forests in Africa and deserts in central Asia. Dormouse species vary significantly in both cranial and mandibular morphology, with some of these morphological variations thought to be related to adaptations to specific habitats and diet (Wahlert et al., 1993; Hennekam et al., 2020a). The fossil record shows the radiation of Gliridae to start during the Early Eocene (Vianey-Liaud, 1994), with the opening of the North Atlantic, and to peak around the late Early Miocene (Daams & de Bruijn, 1995). During this time, dormice were especially abundant and diverse within the Mediterranean area (Hartenberger, 1994), occupying a variety of different ecological niches and exhibiting a great diversity in diet, as evidenced by, for example, the evolution of a hypsodont species (Freudenthal & Martin-Suarez, 2013). The extant dormouse distribution is only a remnant of the success of this family in the past; nonetheless, the various ecological niches occupied by these species today suggest that adaptability to different environments is still retained within the family.

Morphological diversification within animals has been thoroughly investigated in an evolutionary context (e.g. Cardini, 2003; Cardini et al., 2004, 2005; Monteiro et al., 2005; Caumul & Polly, 2005; Michaux et al., 2008). However, these studies often only describe morphological variation at a species or genus level (e.g. Cardini et al., 2004; Ferreira-Cardoso et al., 2020). Morphometrics at a family level sometimes requires the analysis of large numbers of specimens to represent the full extent of this taxonomic rank. Despite some families containing only a small number of species, e.g. Castoridae (2 species), Pedetidae (2) and Aplodontiidae (1), many rodent families are quite species-rich (Burgin et al., 2018). Thus, often a subset of these large families is analysed in order to make the study more comprehensible, as has been done, for example, in the case of Sciuridae (Michaux et al., 2008; Lu et al., 2014). A large number of specimens can often necessitate the use of 2D morphometrics, in which the lack of the third dimension can have significant impact on the strength of the conclusions that can be drawn (Roth, 1993; Álvarez & Perez, 2013; Cardini, 2014; Bakkes, 2017). In particular, more subtle (intra-specific) morphological variation is affected by the inaccuracies seen in 2D datasets (Cardini & Chiappelli, 2020). Fortunately, the increased affordability of 3D scanning of osteological material has a clear positive impact on morphometric studies of large 3D datasets (e.g. Marcy et al., 2018, Ferreira-Cardoso et al., 2020).

Gliridae is a relatively small family within Rodentia with only 29 species in 9 genera (Holden-Musser et al., 2016), making it a tractable group for analyzing morphological variations with respect to size, phylogeny and ecology at this taxonomic level. Previous studies have analysed the cranium within extant dormice (Wahlert et al., 1993; Daams & de Bruijn, 1995; Koenigswald, 1995; Potapova, 2001; Hautier et al., 2008; Hennekam et al., 2020a) and found large variation in, for example, the zygomasseteric construction, morphology of the auditory bullae, and dental characteristics. In particular, the subfamily Graphiurinae appears to differ significantly from other dormice, resulting in much debate on its phylogenetic position within the family (Winge, 1941; Simpson, 1945; Vianey-Liaud & Jaeger, 1996). Molecular phylogenetic studies have not been conclusive for Gliridae, resulting in the positioning of some genera and species remaining unresolved (Bentz & Montgelard, 1999; Montgelard et al., 2003; Nunome et al., 2007).

## Aims

The considerable morphological variation, but relatively low number of species within Gliridae mas it an excellent candidate for a case study investigating adaptive morphological features within a group at species, genus and (sub)family level. Using geometric morphometrics, this study aims to characterise the morphological variation present in the crania and mandibles of dormice and to assess the relationship between morphology and ecology. All but one of the genera comprising Gliridae are represented in the study, resulting in a nearly complete morphological overview of this family. The following three hypotheses will be addressed:

### Intrageneric morphological variation within dormice is less pronounced than intergeneric variation

Clear morphological variation is present both between (Wahlert et al., 1993; Hennekam et al., 2020a), and within dormouse genera (Storch, 1978 and Figure 1). In order to understand the morphological traits within this family, the extent of this inter- and intrageneric morphological variation will be analysed. Given that certain genera are highly specialized to specific habitats (e.g. the desert dormouse *Selevinia*), traits that vary between genera are expected. However, intrageneric morphological variation is also expected for specimens occupying various ecoregions. By evaluating the morphological variation associated with various ecoregions within genera, it is possible to identify to what extent shape variation within Gliridae is the result of phylogenetic divergence, and which morphologies might indicate adaptative features to specific habitats and lifestyles.

**Figure 1.**
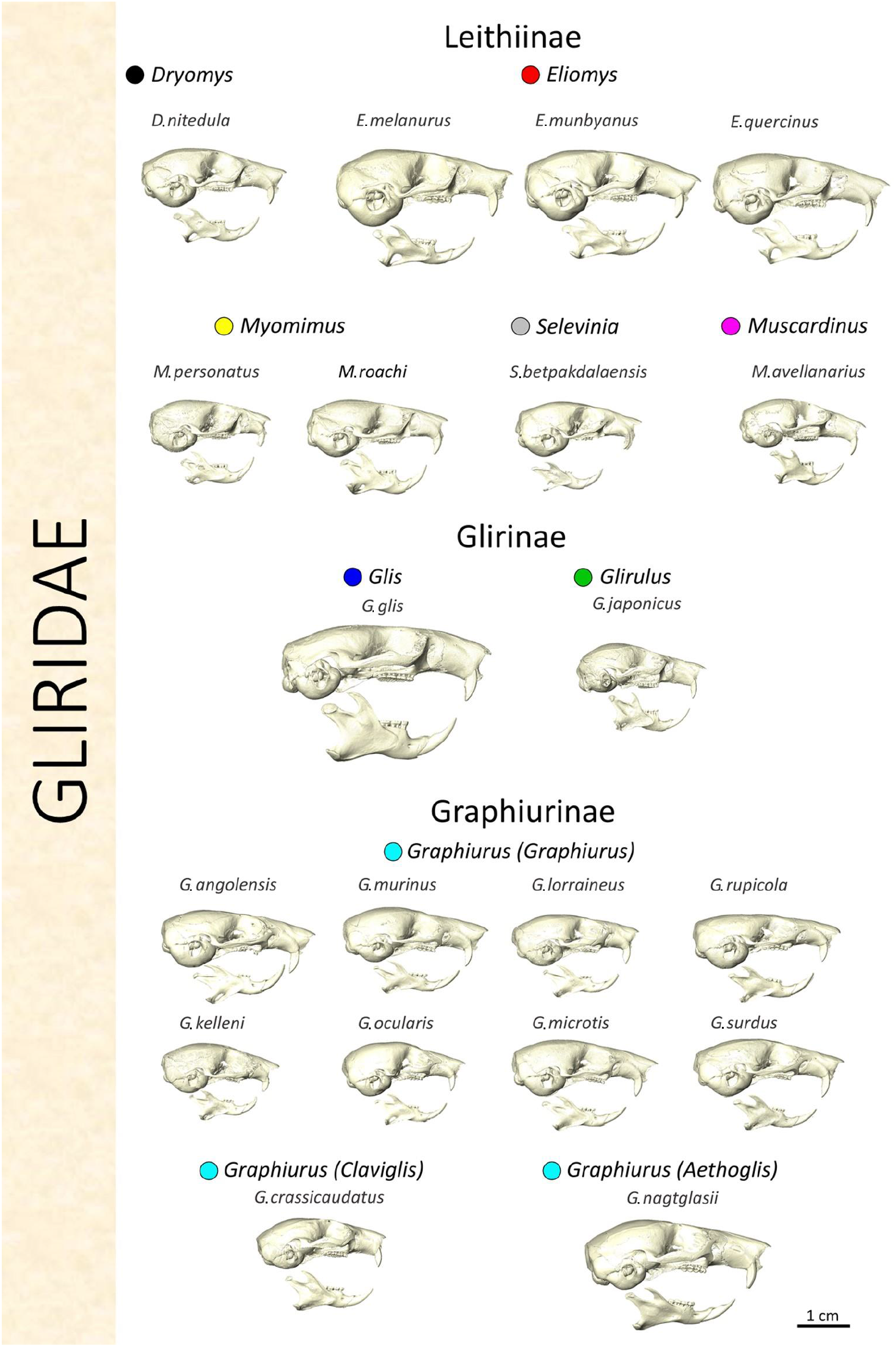
Lateral view of crania and mandibles representing all the species used in this study. Colours are associated to datapoints in subsequent figures referring to the genus. The same scale is used for all specimen.

### Size related variation will have the same direction within and between species

Considering the large size range within Gliridae, a common allometric trend within the family is to be expected. Strong correlations between size and shape have been identified within dormice species (Hennekam et al., 2020b), resulting in species- or genus-specific allometric trajectories. The size distribution within certain genera (e.g. *Eliomys* and *Graphiurus*) is relatively large, covering a substantial proportion of total size variation present within Gliridae. Furthermore, clear size groups are noticeable between genera, indicating size to be linked with speciation and adaptation. Analysing the effects of size on shape at a family level will enable the identification of cranial and mandibular adaptations for coping with increasing body size.

### Correlations between cranio-mandibular morphology and ecological traits/habitat will be significant both within genera and across the entire family

Terrestrial ecoregions are highly dependent on climatological variables. As morphological variations are expected to be linked with various ecoregions (e.g. enlarged auditory bullae in desert-like environments [Hennekam et al., 2020a]), a correlation between climate and morphology is anticipated. Within-genus shape variation will be analysed to identify potential recurring structural adaptations present at a family level.

This study aims to analyse the three-dimensional cranial and mandibular form amongst all dormice. The results of these analyses will then be used to explain size and ecological variables features present at a species/genus level, thus providing a better understanding of the overall morphological variation present within Gliridae.

## Material and methods

### Sample and data acquisition

A total of 109 specimens were included in this research, encompassing 8 of the 9 extant dormouse genera (Figure 1). The missing genus, *Chaetocauda*, is incredibly rare, being known from only five specimens (Holden-Musser et al., 2016). Multiple populations were included for widely dispersed species. Insular populations were avoided, as morphological variation in dormice linked with insularity is described in a previous study (Hennekam et al., 2020b). Even though sexual dimorphism is not considered to be present within dormice (Holden-Musser et al., 2016), whenever possible, an equal representation of both sexes was included within the dataset. The material used is located in the collections of Muséum d’Histoire Naturelle, Paris (MNHN); the Natural History Museum, London (NHMUK); the Senckenberg-Forschungsinstitut und Naturmuseum, Frankfurt (SMF); and the Zoological Institute in St Petersburg (ZISP). Appendix Table S1 gives a detailed list of all specimens. All material was digitized using micro-computed tomography (μCT), using facilities at the X-ray Tomography Facility, University of Bristol (Nikon T H 225 ST CT scanner) and University of Liverpool (SKYSCAN 1272), as well as one scan at the Centre for X-ray Diffraction Studies of Saint Petersburg State University (SKYSCAN 1172). Isometric voxel dimensions ranged between 4 and 40 μm, depending on the size of the specimen and the scanning facility used. As this project focuses on static and evolutionary adaptations related to cranial morphology, only adult specimens (fully erupted third molar) were included to exclude ontogenetic variation from the dataset.

### Data collection

3D anatomical landmarks were placed on surface reconstructions derived from the μCT scans, in order to represent cranial and mandibular shape. Landmarks were recorded in the imaging software Avizo Lite v9.2.0 (Thermo Fisher Scientific, Waltham, MA) using configurations only including type 1 and 2 landmarks (Bookstein, 1991). The cranial landmark set consisted of 42 anatomical landmarks, and the mandibular configuration included 19 landmarks (Figure S1 and S2). Only the left-hand side of the specimens was landmarked. Whenever this area was incomplete or clearly deformed, the object was mirrored digitally to enable landmarking of the right-hand side instead, assuming bilateral symmetry within all dormice.

### Analyses

This study analyses the shape and size of various dormouse species at a family, genus and species level. The initial analyses included all specimens within the family, not correcting for any phylogenetic variation. Subsequently, statistical analyses were performed at a genus level and eventually qualitative analyses at a species (subspecies) level.

All statistical analyses were undertaken in R Studio v3.5.3 (RStudio, Inc., Boston, MA) using various R packages for data preparation and visualisation (*Morpho*: Schlager, 2017; *geomorph*: Adams et al., 2018; *Arothron*: Profico et al., 2018). All configurations were superimposed using a Generalized Procrustes Analysis (GPA) to provide optimal comparability of shape between specimens (Gower, 1975; Rohlf & Slice, 1990). A GPA involves translating all specimens to the origin, optimally rotating using a least-square criterion, and isotropic scaling of all shapes to unit-centroid size in order to align the landmarks as accurately as possible. Centroid size was used as the estimation of size in all subsequent analyses.

### Family level

A principal component analysis (PCA) was used to visualise the shape variance within the entire mandibular and cranial dataset. PCAs including all axes representing over 5% of the total shape variation were plotted. The specimen best approximating the mean shape was warped along the axes to represent the morphological shape variation associated with these components.

Allometry within Gliridae was evaluated using Procrustes ANOVAs for the mandibular and cranial dataset. In order to investigate whether unique (genus-specific) allometries or a common (family-specific) allometric trajectory best explains the correlation between size and shape, ANOVAs including common and unique allometries were performed to identify the correct model for analysing allometry within the datasets.

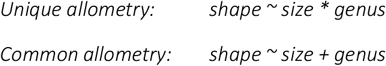

### Genus level

More detailed analyses were performed per genus, applying ANOVAs showing correlations with shape for both size and location. Location is referred from the documented collection site. However, for numerous specimens only the country of origin was documented. For consistency, locations refers to the country of origin of the specimens, unless clear differences in populations were made by the assignment of subspecies (e.g. *Eliomys quercinus lusitanicus*, a subspecies in the South of Spain). These analyses evaluate to what extent intrageneric variation occurs between geographically separated groups, for both size and shape.

### Species level

In order to evaluate the effects of ecological variance on skull morphology within dormice, one genus per subfamily was investigated in more detail, assessing cranial and mandibular shape per geographical region in more detail. Criteria for the chosen genera included a wide dispersal area with specimens collected from various locations, a significant correlation between location and shape, and relative abundance in the dataset. The subfamily Glirinae includes two species; *Glirulus japonicus* and *Glis glis*. The former is only known from Japan, whereas *Glis glis* is widely distributed across Europe and small parts of Asia. Within Leithiinae, *Eliomys* is considered the most widely dispersed genus and is considerably more abundant in the dataset than other genera belonging to this subfamily. Graphiurinae is a monogeneric subfamily, resulting in *Graphiurus* being the only potential candidate, which fortunately meets all the determined criteria. Thus, the genera *Glis, Eliomys* and *Graphiurus* were considered the best candidates for analysing shape variance with respect to various habitats. A relatively large number of specimens of these genera were included in the datasets (12, 39 and 30 respectively) and all three genera are widely distributed; together, they cover the majority of the total dispersal area of all dormice, excluding the eastern parts of Asia. The method of analysing shape is dependent on the number of specimens within the genus, the species included and how they morphologically differ, and the different ecoregions occupied. Variations in morphology were quantified by comparing species to the mean Procrustes shape per genus. Only a limited number of specimens per location were analysed, as to avoid overrepresentation affecting the mean shape.

### Validation and caveats

For the Eurasian specimens, the assignment of location is often not linked with a change in ecoregion. Instead, physical barriers (e.g. rivers, lakes and mountain ranges) are more likely to play a role in the segregation of populations. The Sub-Saharan ecoregions are more variable and seem to act as physical barriers as well. However, the species-rich genus *Graphiurus* is the only dormouse genus inhabiting this part of the world. The dispersal of species within this genus is clearly associated with climatic variables, such as temperature and precipitation, and the associated ecoregions (Appendix Figures S3 and S4). Unfortunately, the *Graphiurus* dataset is limited, and does not include all species. Furthermore, for some species only one specimen is included, resulting in a poor representation of the species. Lastly, the lack of precise geographical information on some specimens resulted in an inaccurate estimation of climatic variables and in some cases in the assignment of two different ecoregions. Considering the large number of different species and the wide distribution of certain species within the genus *Graphiurus*, covering a large number of ecoregions, it is likely that this study has captured only a fraction of the morphological variation present. It must be noted that ecoregions were not used in statistical analyses, but instead were assessed qualitatively.

## Results

### Shape variation (Family level)

The distribution of dormice in shape space is shown in the PCA plots in Figures 2 and 3 and described below. A consensus skull (MNHN 1961-534; *E. quercinus*) and mandible (NHM 61.362; *D. nitedula*) were warped along the principal components in order to visualise the shape variation present within Gliridae (Figures 2 and 3). Tables 1 and 2 describe the larger shape variations occurring on these axes, as well as the groups clearly associated with these traits. The skulls and mandibles of specimens representing all the species analysed in this study are displayed in Figure 1. The species are colour-coded in correspondence with the datapoints in subsequent figures.

**Table 1:**
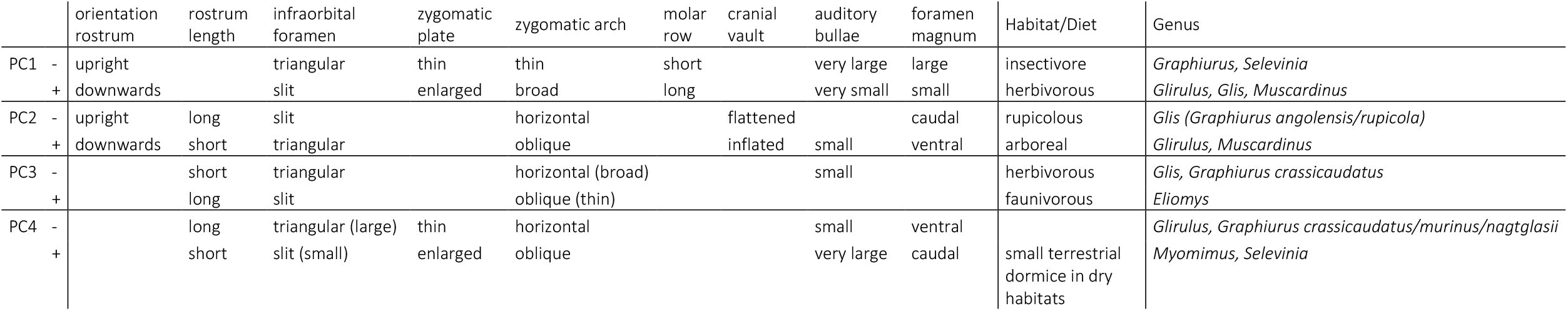
Description of the shape variation occurring on the first four principal component axes for the cranial data set and references to species associated with more positive or negative values.

**Table 2:**
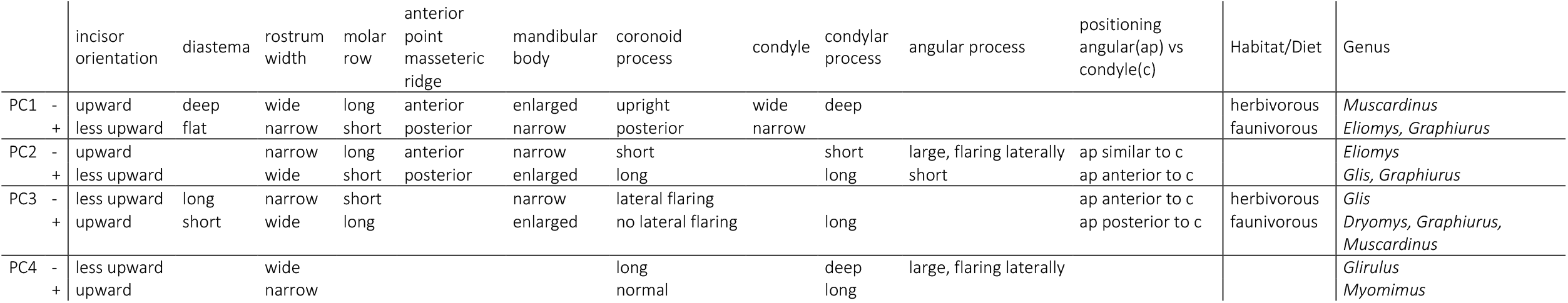
Description of shape variation occurring on the first four principal component axes for the mandibular data set and references to species associated with more positive or negative values.

**Figure 2:**
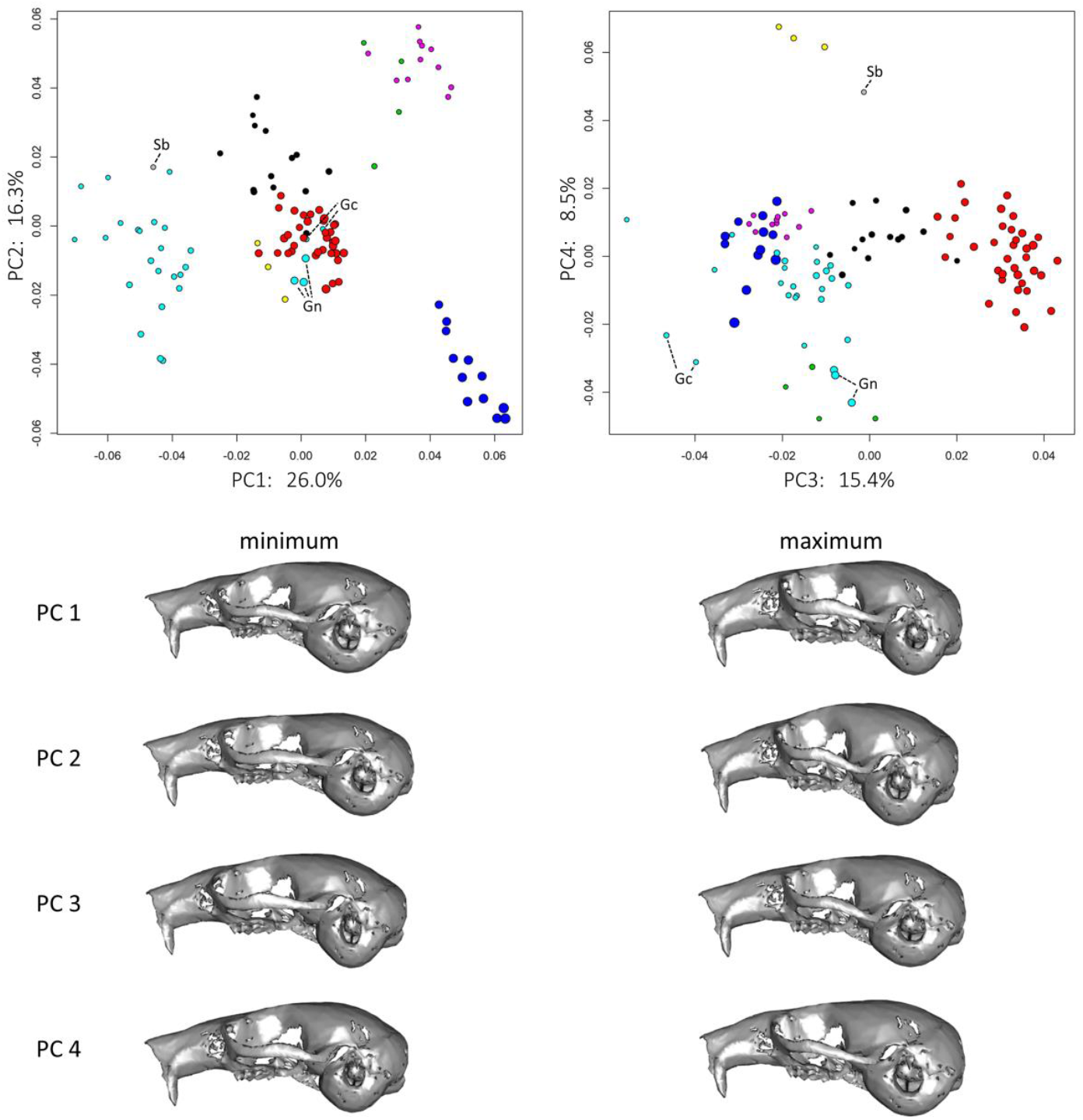
*PCA for the cranial dataset and corresponding warps on the first four principal components. The skull of an* Eliomys quercinus *specimen from France (MNHN: 1961-534) is most similar to the mean cranial shape in the dataset and used for the warping along the axes. Gc =* Graphiurus crassicaudatus; *Gn =* Graphiurus nagtglasii; *Sb =* Selevinia betpakdalaensis. *Colour key (also see Figure 1): cyan =* Graphiurus; *grey =* Selevinia; *black =* Dryomys; *red =* Eliomys; *yellow =* Myomimus; *purple =* Muscardinus; *green =* Glirulus; *blue =* Glis.

**Figure 3:**
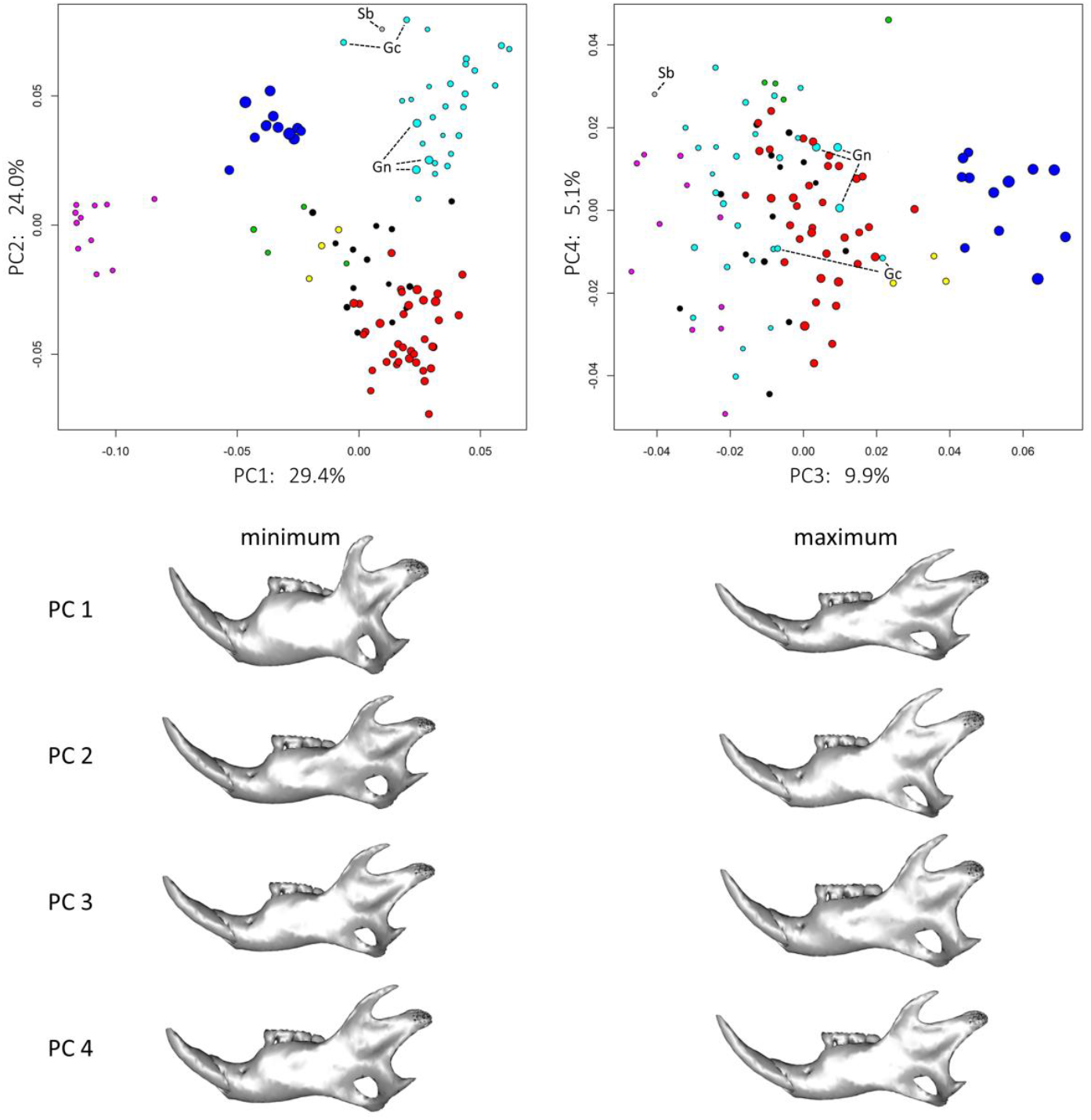
*PCA for the mandibular dataset and corresponding warps on the first four principal components. The mandible of an* Dryomys nitedula *specimen from Turkey (NHMUK: 61*.*362) is most similar to the mean mandibular shape in the dataset and used for the warping along the axes. Gc =* Graphiurus crassicaudatus; *Gn =* Graphiurus nagtglasii; *Sb =* Selevinia betpakdalaensis. *Colour key: cyan =* Graphiurus; *grey =* Selevinia; *black =* Dryomys; *red =* Eliomys; *yellow =* Myomimus; *purple =* Muscardinus; *green =* Glirulus; *blue =* Glis.

#### Crania

The first 32 principal component axes explain 95% of the total shape variance within the cranial dataset. The first four axes are the only components that explain more than 5% of the total variance and are plotted in Figure 2. Clear morphological variation between genera is apparent in these four components, with every genus being separated from another in at least one of the axes. The first principal component (PC1) does not show a clear size signal, but instead distinguishes between the three subfamilies; Graphiurinae for negative values, Leithiinae in the centre and Glirinae at more positive values. Some specimens deviate from this pattern, as two species within *Graphiurus* (*G. nagtglasii* and *G. crassicaudatus*) seem to be more similar to Leithiinae specimens, whereas *Selevinia* is placed within the Graphiurinae on the first principal component. *Muscardinus*, previously assigned to Glirinae but currently considered part of Leithiinae, shows relatively positive values on the first axis. For the first three principal components, the cranial morphology of *Muscardinus* appears to resemble that of the Japanese dormouse *Glirulus japonicus*, placed within the Glirinae. The second principal component differentiates between genera within sub-families, as clear separation visible between *Glis glis* (negative) and *Glirulus japonicus* (positive) for Glirinae and between *Eliomys, Dryomys* and *Muscardinus* (negative to positive) in Leithiinae. On this axis larger species are placed at more negative values (*Eliomys, Glis* and certain *Graphiurus* species), and the smaller species at more positive levels (*Muscardinus* and *Glirulus*). The third principal component also shows a clear segregation between species, however, the size element is lost as the two largest genera, *Glis* and *Eliomys*, are at opposite ends of the spectrum. *Myomimus* within Leithiinae is now separated from *Eliomys* and *Dryomys*, and instead resembles the smaller dormice *Glirulus* and *Muscardinus*, as well as some *Graphiurus* specimens. The fourth principal component indicates morphological similarities between *Myomimus* and the desert dormouse *Selevinia*. Furthermore, the Japanese *Glirulus* resembles certain *Graphiurus* species (*G. crassicaudatus, G. nagtglasii* and some *G. murinus*) at the more negative values on this axis.

#### Mandibles

For the mandibular principal component analyses, 21 axes are needed to explain 95% of the total shape variation, with the first four components explaining more than 5% of shape variation each (Figure 3). Clear separation between genera is visible within these axes, whereas size does not appear to be a prominent driver of shape variation. The first principal component separates *Muscardinus* from all other genera, which more closely resembles Glirinae genera in shape, than the Leithiinae, to which it supposedly belongs. The second principal component distinguishes between Graphiurinae and Leithiinae. In contrast to the cranial dataset, the mandibular shape of *G. nagtglasii* and *G. crassicaudatus* places them neatly within their subfamily. The desert dormouse *Selevinia* is still located within the Graphiurinae, whereas it is actually thought to belong to Leithiinae. *Glis* is associated with more positive values on principal component 3. Furthermore, *G. crassicaudatus* and *G. nagtglasii* show more positive values than other *Graphiurus* species, a trend also seen in the cranial analysis for these two species (Figure 2: PC1 and PC4). The fourth principal component for the mandibular shape variation is less indicative for morphological variation between genera in comparison with the fourth component in the cranial dataset, albeit still clustering species within genera such as *Glirulus* and *Myomimus*.

### Size variation (Family level)

Based on the landmark configuration, the centroid size of each dormouse genus was determined for both skull and mandible (Figure 5). Clear size variation is present between genera, with the largest dormouse being *Glis glis* (centroid size = 93), and the smallest dormouse being *Muscardinus avellanarius* (46). These size differences are even more pronounced within the mandibular dataset (min = 18, *Graphiurus murinus*; max = 43, *Glis glis*). *Graphiurus*, being by far the most species-rich dormouse genus, showed the largest intrageneric variation in centroid size, with the largest skull being 56% larger than the smallest specimen, and the largest mandible being 69% larger. This size variation within *Graphiurus* is the result of the exceptionally large species G. *nagtglasii* (outlier in Figure 4). It should be noted that due to the scarcity of certain dormouse species, some genera are underrepresented. The maximum within-genus size difference is therefore expected to be larger than displayed in Figure 4. This absence within the dataset is exceptionally notable for the genus *Eliomys*, for which insular specimens were excluded from this study, but the largest population of which is found on the island of Formentera (Storch, 1978; Hennekam et al., 2020b). The results indicate that relatively large dormouse genera (*Eliomys* and *Glis*) are more variable in size than smaller genera (*Muscardinus* and *Glirulus*). This can partly be explained by the large sampling of *Eliomys* (39 specimens) and the poor sampling for *Glirulus* (4 specimens). Nonetheless, *Muscardinus* and *Glis* are relatively equally represented in the dataset (11 and 12 specimens respectively), with size variation in the larger *Glis* species being clearly more pronounced.

**Figure 4:**
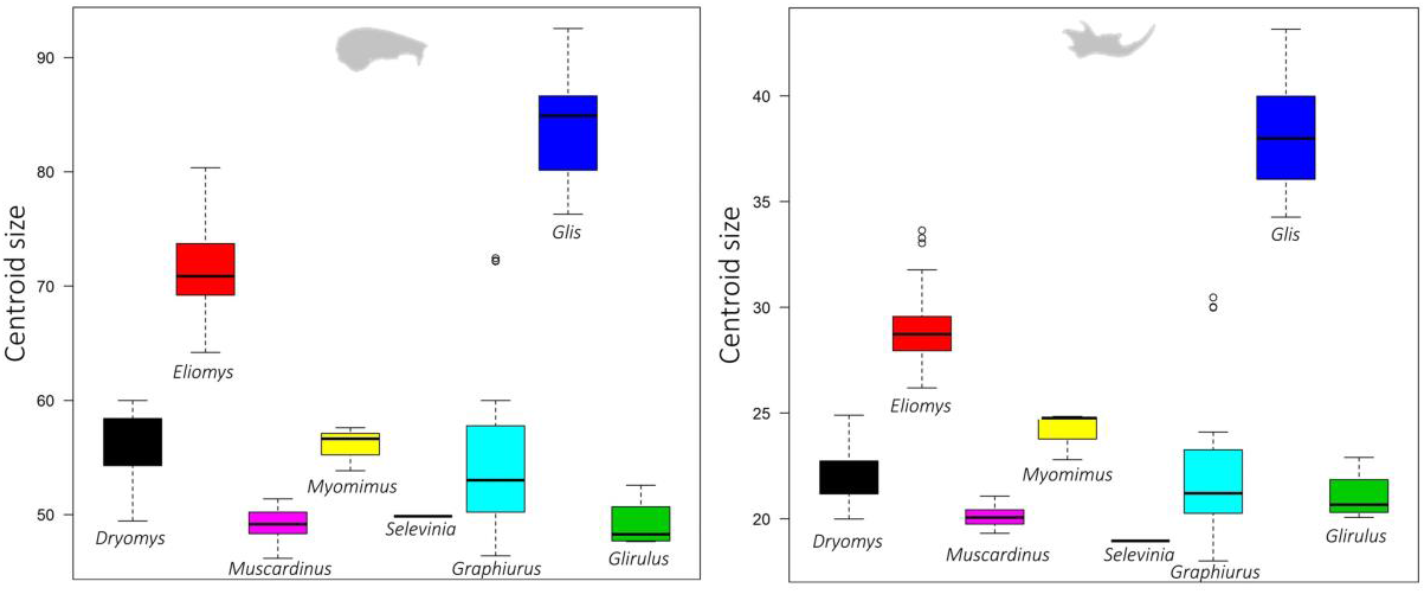
Size variation between dormouse genera in both the cranial (left) and mandibular (right) dataset.

**Figure 5:**
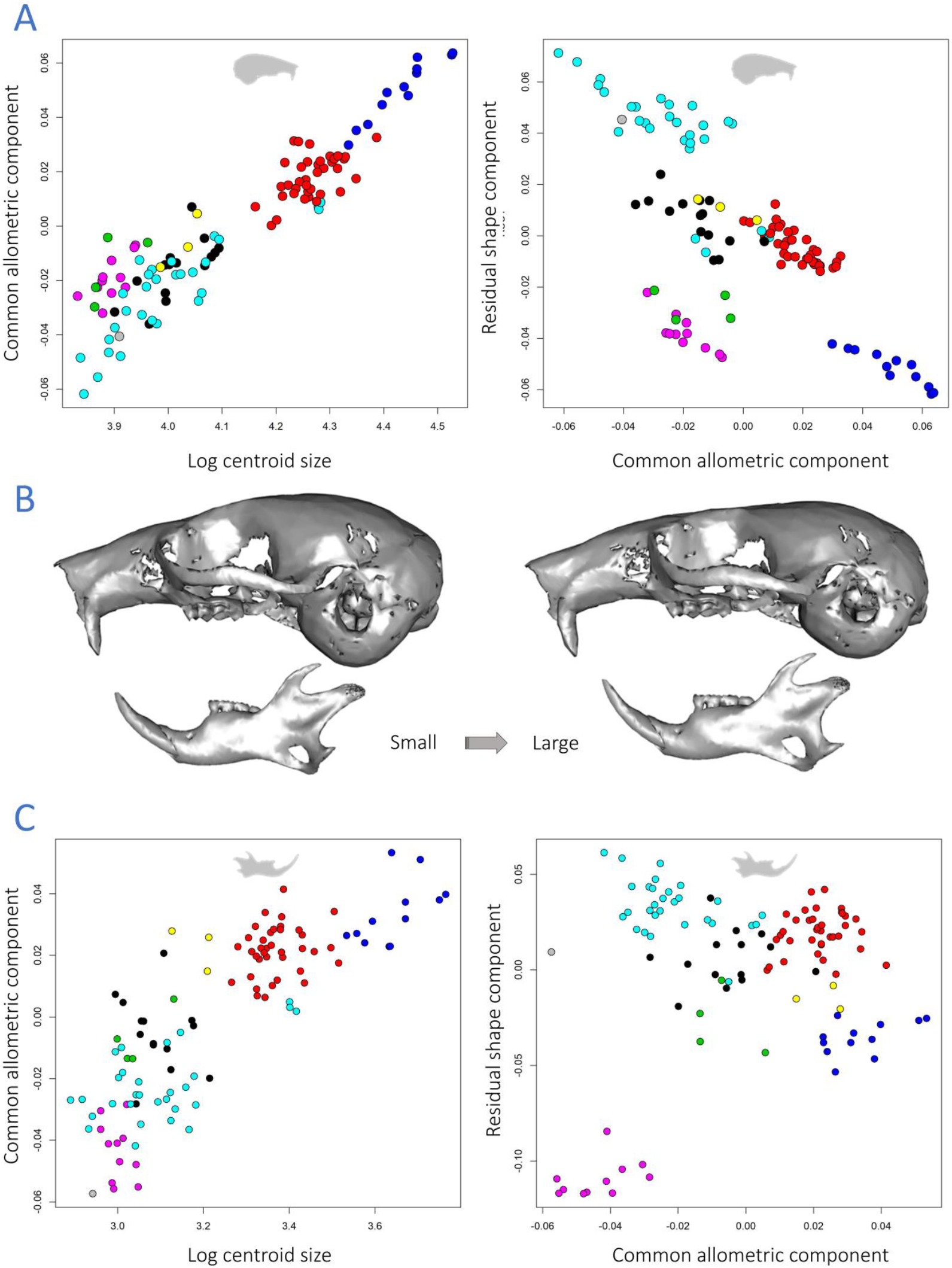
*Common allometric component analyses for the cranial (A) and mandibular (C) dataset, including corresponding warps (B) representing the morphology of smaller dormice (left) and larger specimens (right). Colour key (also see Figure 1): cyan =* Graphiurus; *grey =* Selevinia; *black =* Dryomys; *red =* Eliomys; *yellow =* Myomimus; *purple =* Muscardinus; *green =* Glirulus; *blue =* Glis.

### Allometry (Family level)

The model comparisons between the unique allometries (shape ∼ size * genus) and the common allometries (shape ∼ size + genus) were not significant. These results indicate the common allometric component (without grouping) to be the most accurate model for analysing allometry within dormice, rather than using unique allometric trajectories per genus (Figure 5, and see Appendix Table S2 for test). In total, 7.5% of the shape variance in the mandibular dataset was explained by size, with 16.7% explaining the variance in cranial shape. These relatively low values clarify why the effects of size on shape are not well represented in the first principal components (Figures 2 and 3). The linear relationship between size and shape was tested using a Pearson product-moment correlation coefficient, showing strong correlations in both datasets (cranium: r = 0.934, 95% CI = 0.91-0.95; mandible: r = 0.829, 95% CI = 0.76-0.88). The amount of shape variance is greater and the linear relationship between shape and size is considerably stronger in the cranial dataset, indicating that allometry influences the morphology of cranial features to a greater degree than in the mandible.

### Cranial allometric variation

Smaller individuals are characterised by a short and robust rostrum and a more bulbous cranial vault (Figure 5). Larger specimens display a more elongated snout and a clear flattening of the skull. The infraorbital foramen is more triangular in shape in smaller dormice, whereas it is represented by an elongated opening in large dormice. Furthermore, the auditory bulla appears to be relatively small in larger specimens. Lastly, the foramen magnum is large and ventrally orientated in small dormice, but relatively smaller and more posteriorly orientated as size increases.

### Mandibular allometric variation

With increasing size, the angular process in the mandible moves slightly posteriorly as the condylar process moves more anteriorly, resulting in the posterior margin of both processes to align with each other (Figure 5). Furthermore, there is a small decrease in the relative length of the mandibular diastema in larger specimens.

### Shape and size variation (Genus level)

Procrustes ANOVAs per genus resulted in significant correlations between shape and independent variables size and location for some genera, but not in others (Table 3; Appendix Tables in S3). For the genera *Selevinia, Myomimus* and *Glirulus*, only specimens originating from the same location were included in the dataset. As no variable for locality could be included, size was used as the single independent variable for the analysis. In addition, these genera were relatively underrepresented within the dataset (1, 3 and 4 specimens respectively), therefore, the correlation between size and shape could not be assessed. *Muscardinus* showed a significant correlation for both cranial and mandibular shape with location, but not with size. The same trend was observed in the genus *Dryomys*, with location being significant in both instances, even though size was not significant for the mandibular dataset and only just significant for the cranial dataset (p = 0.041). The larger dormouse genera *Glis* and *Eliomys* showed significant correlations for both location and size on shape, as well as significant interaction terms. The species-rich genus *Graphiurus* indicated significant shape correlations as well in both datasets, although the interaction term between location and size for the mandibular term was not significant.

**Table 3:** Procrustes ANOVAs per dormouse genus for both cranial (top) and mandibular (bottom) datasets, evaluating the shape variation with respect to centroid size and location: shape ∼ size * location. Relevant and statistically significant values are bold, with the p-values summarised with asterisks (p = 0.01 -0.05, * ; p = 0.001 -0.01, **). NA, not applicable (only one specimen and/or location included).

| subfamily | Genus | # specimens | R<sup>2</sup> size | R<sup>2</sup> location | R<sup>2</sup> interaction |
| --- | --- | --- | --- | --- | --- |
| Leithiinae | *Dryomys* | 15 | **0.11** * | **0.52** * | **0.13** ** |
|  | *Eliomys* | 35 | **0.06** ** | **0.48** ** | **0.19** ** |
|  | *Muscardinus* | 11 | 0.11 | **0.46** ** | 0.09 * |
|  | *Myomimus* | 3 | 0.68 | NA |  |
|  | *Selevinia* | 1 | NA | NA |  |
| Glirinae | *Glirulus* | 4 | 0.49 | NA |  |
|  | *Glis* | 12 | **0.28** ** | **0.60** ** | **0.04** ** |
| Graphiurinae | *Graphiurus* | 30 | **0.17** ** | **0.39** ** | **0.1** * |

| subfamily | Genus | # specimens | R<sup>2</sup> size | R<sup>2</sup> location | R<sup>2</sup> interaction |
| --- | --- | --- | --- | --- | --- |
| Leithiinae | *Dryomys* | 15 | 0.07 | **0.63** ** | 0.11 |
|  | *Eliomys* | 37 | **0.05** * | **0.50** ** | **0.13** ** |
|  | *Muscardinus* | 11 | 0.15 | **0.44** ** | 0.05 * |
|  | *Myomimus* | 3 | 0.56 | NA |  |
|  | *Selevinia* | 1 | NA | NA |  |
| Glirinae | *Glirulus* | 4 | 0.48 | NA |  |
|  | *Glis* | 12 | **0.18** ** | **0.72** ** | **0.04** ** |
| Graphiurinae | *Graphiurus* | 30 | **0.06** ** | **0.41** ** | 0.1 |

### Shape and size variation (Species level)

*Eliomys, Glis* and *Graphiurus* were evaluated as representatives of their associated subfamilies (Leithiinae, Glirinae and Graphiurinae, respectively). Shape variation within all three genera is substantial, albeit that some genera are more speciose than others. Depending on the genus, the number of specimens within the dataset and the various locations represented, different methods for assessing within-group shape variation based on GPAs and PCAs were applied. The descriptions of the monospecific genus *Glis* was focused on the most distinctive specimens within the dataset. The ten most distinctive specimens were described for the genus *Eliomys*. This study follows Pavlinov & Potapova (2003) by splitting the genus *Graphiurus* into three subgenera. These are: *Aethoglis*, which contains *G. nagtglasii*; *Claviglis*, containing *G. crassicaudatus*; and *Graphiurus*, which contains all other *Graphiurus* species. The subgenera *Graphiurus* (*Aethoglis*) and *Graphiurus* (*Claviglis*) are morphologically very distinct from the subgenus *Graphiurus* (*Graphiurus*). Their skull morphology was described individually, whereas a GPA and PCA was used to determine the ten most distinct *Graphiurus* (*Graphiurus*) specimens.

### Glirininae: Glis glis

This species shows a significant correlation between shape and size, as well as shape and location, including the interaction term (Table 3). The edible dormouse *Glis glis* is the only extant species in the genus *Glis* and is considered the largest extant dormouse species (Storch, 1978). The distribution of the species is mainly characterized by temperate broadleaf, coniferous and mixed forests throughout Europe and southwestern parts of Asia.

Specimens representing populations from seven different localities across Eurasia were analysed based on shape, size and ecoregion. The genus is represented by 12 individual specimens within the dataset, resulting in a relatively low number of specimens per locality. It is therefore difficult to assess whether these individuals are representative for their population. The skull and mandible size of *G. glis* is quite variable within populations, especially with respect to relatively small to medium sized specimens (centroid skull size: 76.3 -86.7). Dormice from Azerbaijan, Italy and Germany show similar skull sizes (86.6 -86.7), although climate varies significantly between these localities. The second-largest *Glis* specimen is significantly larger than the rest of the mainland dormice (92.4) and belongs to the subspecies *G. g. persicus*, originates from Iran and occupying a desert-like habitat (Figure 6).

**Figure 6:**
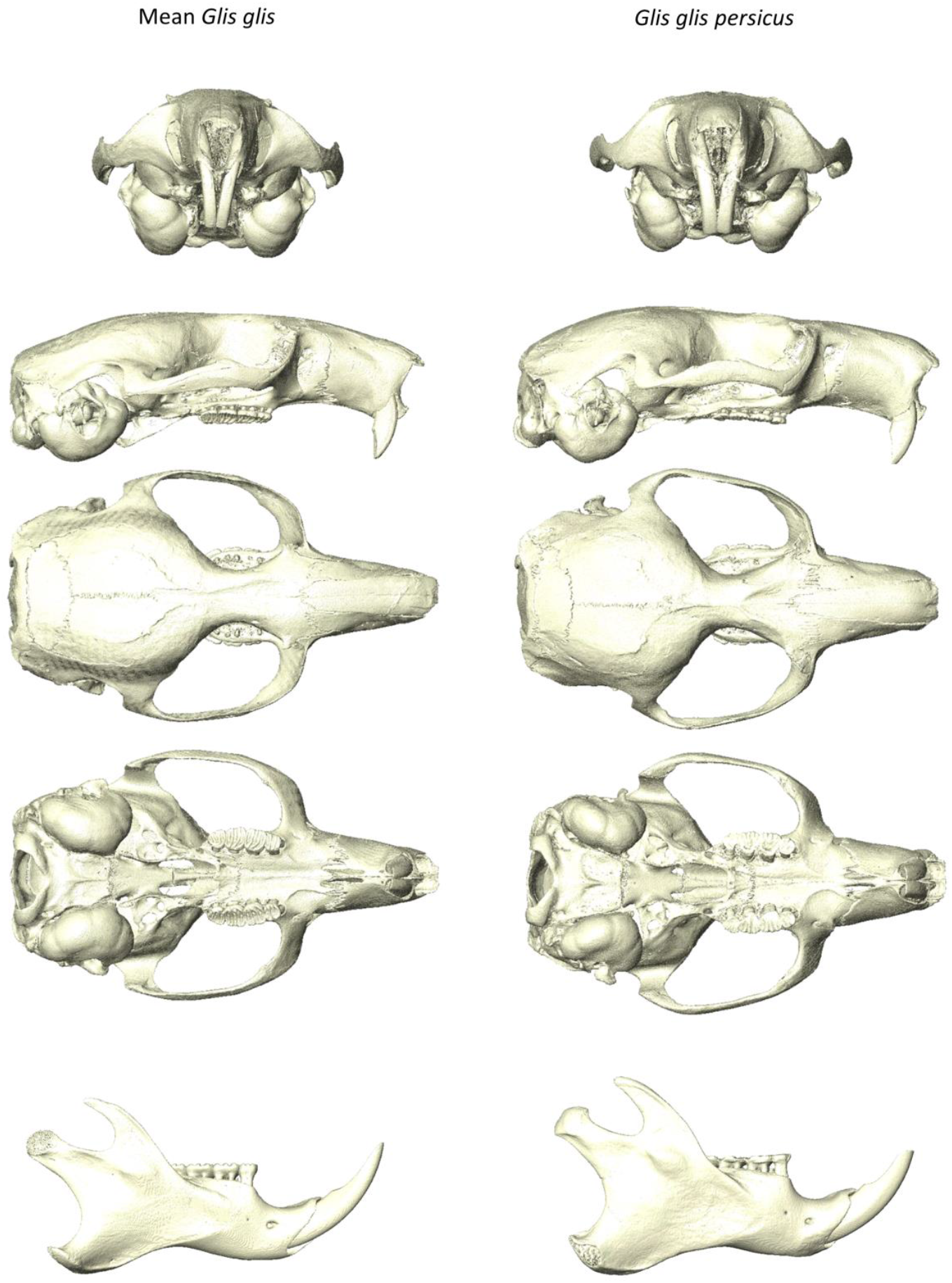
*Mean crania (SMF: 82070) and mandible (SMF: 40073) of* Glis glis *on the left, and the distinctive* Glis glis persicus *(NHMUK: 27*.*10*.*26*.*21) from a frontal, lateral, dorsal and ventral perspective on the right*.

The largest *Glis glis* is found on the island of Sicily (92.5). Due to the tendency of dormice to become larger in isolated habitats, insular dormice were initially excluded from this study. However, the relative scarcity of *Glis glis* within the dataset justified the inclusion of this particular specimen. Interestingly, the Sicilian dormouse is reported to have the smallest body size among all Italian populations (Milazzo et al., 2003), which does not correspond with the findings here, nor the observations by Storch (1978). The skull of this specimen is considerably flatter compared to skulls in other populations. The largest Procrustes distance to the mean is present in the Iranian dormouse (0.0540 for the skull, 0.0559 for the mandible), which is the only specimen inhabiting a desert environment. Morphologically it is most similar to the nearby Azerbaijan population with both populations having a relatively broad rostrum. The nasal bone in the Iranian dormouse flares more laterally anteriorly compared to other populations. Furthermore, the orbit is enlarged and the midorbital constriction is narrower in this specimen, resulting in the fusion of the cranial crests, whereas they are separated in all other populations. When orientated ventrally, the muscle attachment site of the superficial masseter is expanded medially, which is indicative for more strongly developed musculature. The infraorbital foramen in this specimen is elongated superiorly and the zygomatic plate less angled inferoposteriorly. The molars are severely worn, especially lingually, which has resulted in a more ventral orientation of the occlusal surface. The muscle attachment area for the temporalis muscle on the vault is less pronounced. The ventral margin of the masseteric ridge on the mandible is positioned slightly more inferior, effectively enlarging the masseteric region and attachment area for the masseter muscle. Furthermore, the angular process in the Iranian dormouse is more robust.

### Leithiinae: Eliomys

*Eliomys, Dryomys* and *Muscardinus* all show shape variation significantly correlating with location (Table 3). Of the three genera, *Eliomys* is considerably more abundant within the dataset (39, 15 and 11 respectively). Furthermore, this is the only genus with significant correlations between shape and both size and location, as well as the interaction term, for both datasets. The garden dormouse genus *Eliomys* contains three extant species. The species are separated geographically from each other, occupying parts of Northern Africa, Europe and the Middle East. *E. quercinus* inhabits forests across Europe and, in contrast to *Glis glis*, also the Iberian peninsula. *E. munbyanus* occupies the Maghreb region, inhabiting both forest and desert-like environments (Holden-Musser et al., 2016). The patchy distribution of the Asian garden dormouse *E. melanurus* ranges from the north east of Libya to the Middle East, where it occupies very arid environments.

The initial six principal components all explain more than 5% of the total shape variation, which can be an indication that morphological variance within this group is relatively subtle. Segregation between the European *Eliomys quercinus* and the other two species is visible on the first principal component (Figure 7). The cranial morphology of the ten most distinct specimens was evaluated by comparing the landmark positioning with that of the mean shape derived from the GPA. The Procrustes distances from the mean in all specimens varied from 0.043 to 0.027, indicating the overall morphological variation within this genus to be relatively small. Most variation is present in the rostrum, the zygomatic arch and the foramen magnum. *E. quercinus* tends to have an anteriorly displaced its incisive foramen with respect to the other species. Furthermore, the auditory bulla in this species is slightly less inflated. The dorsalmost point of the zygomatic arch in both *E. melanurus* and *E. munbyanus* is positioned more anterosuperiorly. The incisors in *E. munbyanus* appear to be more opisthodont and the foramen magnum is oriented more inferiorly. The largest *Eliomys* specimen in the dataset originates from Sevilla and shows clear flattening of the skull, a feature also seen in the larger *Glis glis* specimens. Within *E. quercinus*, the Swiss populations and, to a lesser extent, the northern Italian populations have the dorsalmost section of the zygomatic arch located more posteriorly compared to all other populations. Furthermore, in the majority of these specimens, the orientation of the foramen magnum appears more caudal.

**Figure 7:**
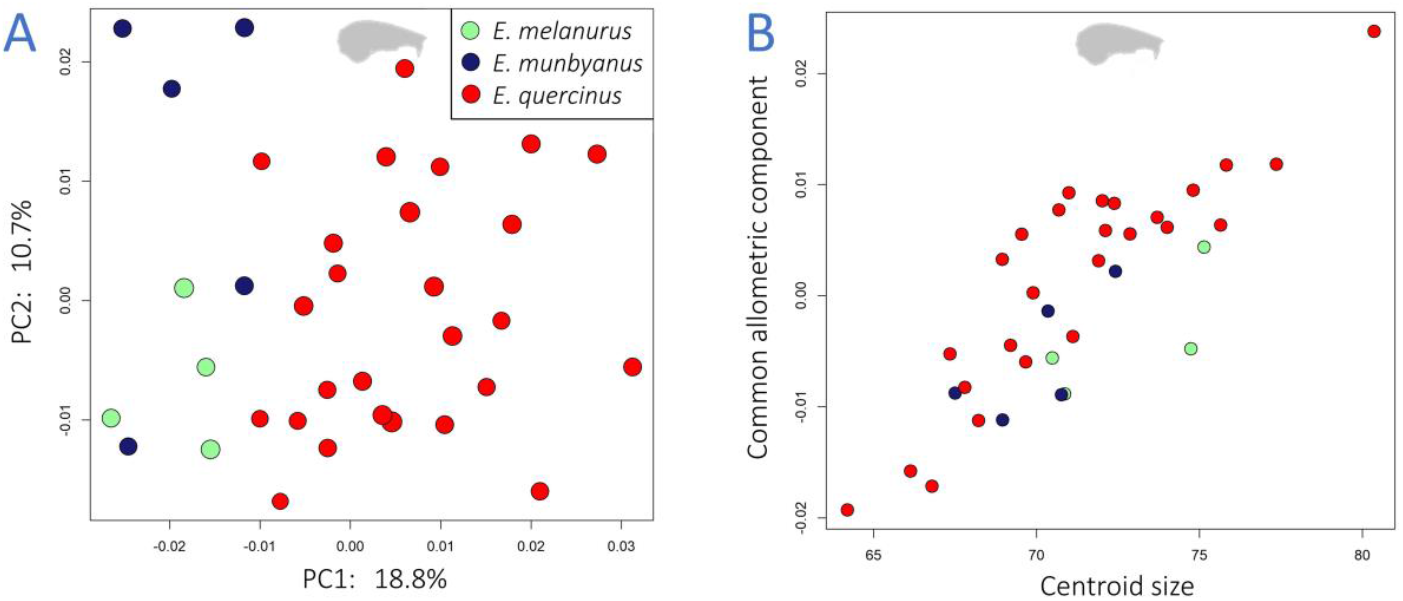
A: PCA of the cranial dataset for the genus Eliomys, displaying the first two principal components describing 29.5% of the total shape variation within this group. B: the common allometric component analysis on the cranial dataset.

The principal component analysis on the mandibular dataset of *Eliomys* also has six principal components explaining more than 5% of the total shape variation. The Procrustes distances are larger than those seen in the cranial dataset, varying from 0.079 to 0.033 from mean. The PCA shows clustering of the Maghreb dormice at more positive values of the first principal component, whereas one specimen of *E. munbyanus* is not placed here (Figure 8A). Two *E. quercinus* mandibles from Sevilla are the largest mandibles within the dataset and clustered with the Maghreb dormice. Furthermore, other relatively large *Eliomys* specimens are associated with positive values on the PC1. The common allometric component analysis indicates that most *E. munbyanus* and *E. melanurus* represent larger *E. quercinus* specimens, whereas not increasing in size themselves (Figure 8B).

**Figure 8:**
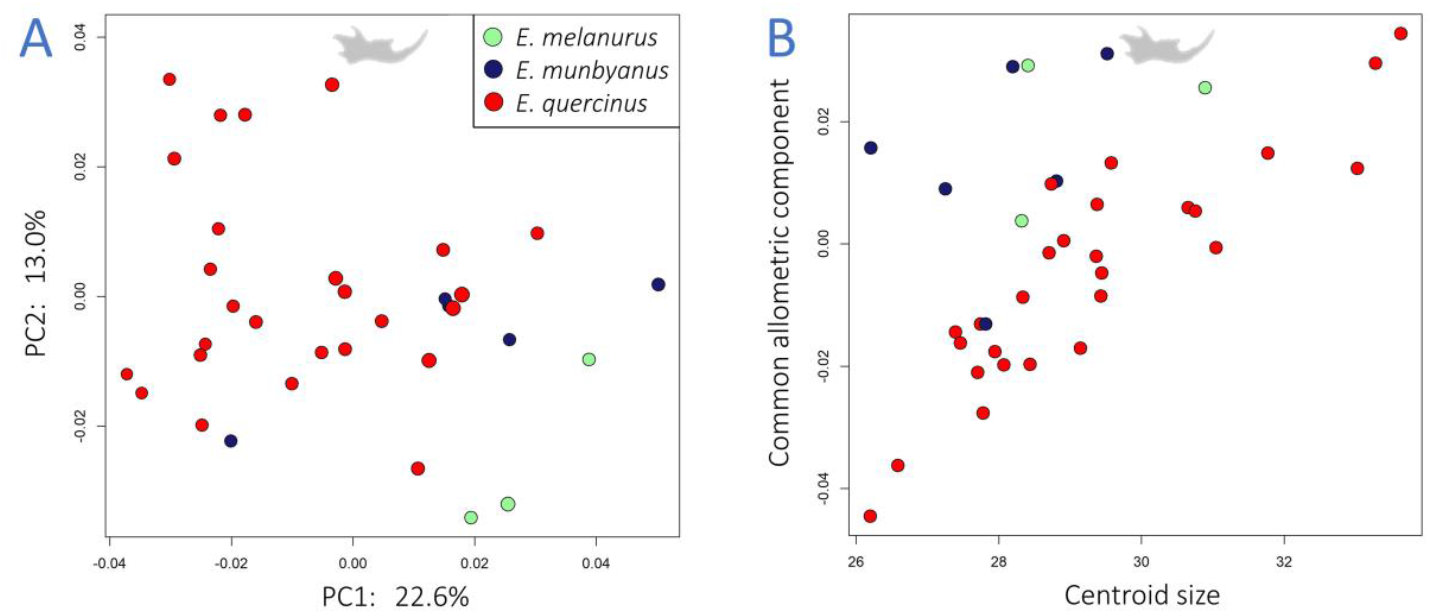
A: PCA of the mandibular dataset for the genus *Eliomys*, displaying the first two principal components describing 35.6% of the total shape variation within this group. B: the common allometric component analysis on the mandibular dataset, indicating that *E. munbyanus* and *E. melanurus* are morphologically similar to larger *E. quercinus* specimens.

### Graphiurinae: Graphiurus

Within the *Graphiurus* subgenus, including a total of 13 different species, significant diversity in cranial morphology is apparent (Figures 1 and 2). The dataset used in this study incorporates eight out of the thirteen extant species. With 14 specimens, the species *G. murinus* is overrepresented in the dataset, but does show clear morphological variation between geographically separated populations (Table 3).

Based on the landmark configuration, the positioning of various cranial elements was analysed. Larger specimens appear to be characterized by a flattening of the vault, a trait seen also within *Glis* and *Eliomys*. Furthermore, enlarged specimens tend to have the foramen magnum positioned more superiorly, resulting in the foramen being oriented more caudally. Smaller specimens are characterised by a shortened rostrum, with the anterior part of the nasal bone deflected more inferiorly. The cranial vault in these specimens is relatively enlarged and more rounded. The auditory bullae are more inflated in specimens occupying dry areas. The rupicolous species *G. rupicola* and *G. ocularis* both show a relatively narrow zygomatic arch and an inflated auditory bulla, whereas only the larger *G. rupicola* has a clearly flattened skull. The *G. ocularis* in this study is rather small, whereas this species is often regarded as the second largest *Graphiurus* species (after *G. nagtglasii*; Holden-Musser et al., 2016). It is unclear if this specimen was labeled incorrectly, or simply represents a relatively small individual. The overrepresented *G. murinus* is known from three localities: Democratic Republic of the Congo, Tanzania and a single specimen from Kenya. The Tanzanian specimens appear to be larger, with the exception of specimens inhabiting ecoregion 7 (Tropical and Subtropical Grasslands, Savannas and Shrublands). The populations from Congo are also assumed to inhabit this ecoregion, whereas the average-sized Tanzanian populations occupy ecoregion 1 (Tropical and Subtropical Moist Broadleaf Forests) and the largest specimens ecoregion 14 (Mangroves). The specimen from Kenya is from a desert environment and is of average size. The cranial morphology of this specimen is very similar to the mean shape of *Graphiurus murinus*. However, the mandibular shape of this particular specimen is very distinctive compared to other *G. murinus* mandibles. Other mandibular and cranial features, including the incisive foramen and the dorsal bending of the zygomatic arch, appear variable within *Graphiurus*, although not distinguishing between species and/or ecoregion.

The monospecific subgenus *Aethoglis* includes the largest extant *Graphiurus* species, *G. nagtglasii*. This specimen is known to be primarily arboreal and rarely seen on the ground (Holden-Musser et al., 2016). Alongside its relatively large size, the skull of *G. nagtglasii* has a large rostrum. The zygomatic arch is relatively robust and the posterior section of the skull appears to be angled inferiorly, resulting a bulbous cranial vault and a more ventral orientation of the foramen magnum. The auditory bullae are relatively small. The molars within this species appear to be less concave and more complex (molar ridges) and robust compared to most other *Graphiurus* specimens. The mandibular shape of this species is not as distinctive as the cranium (Figures 2 and 3).

*G. crassicaudatus* is considered the sole member of the subgenus *Claviglis* and the most herbivorous of all *Graphiurus* species. The zygomatic arch in this species is very pronounced, being more robust than in any other species, including muscle attachment sites on the zygomatic plate that are very well defined. Furthermore, distinct temporal crests are present, as also seen *Glis glis*, although the crest morphology in the two species is different. The orbital constriction in *G. crassicaudatus* is very broad and the rostrum relatively shortened and quite narrow. Lastly, the species shows clear lateral ridges on its robust molars and displays molarised premolars, whereas molars in other *Graphiurus* species tend to be relatively small and concave and lacking distinctive molar ridges. The size of the *G. crassicaudatus* mandible is similar to that of the relatively large *G. angolensis*, whereas the skull of this species is quite small, only slightly larger than the small *G. lorraineus* (see also Figure 1).

## Discussion

Clear morphological variation between and within dormouse populations is present. As well as distinguishing between genera, this study identifies particular changes in shape associated with size (allometry), whereas other morphologies are linked with specific ecological parameters such as habitat and diet.

### Intrageneric versus intergeneric morphological variation

Variability in cranial and mandibular morphology is apparent within Gliridae. With clearly separated genera in shape space, it is evident that the intergeneric variation across the whole family is more pronounced compared to shape variation within genera. However, intrageneric variation in morphology is present and sometimes exceeds intergeneric variation. This is the case for cranial morphology in two subgenera within *Graphiurus, Claviglis* and *Aethoglis*, which are instead more similar in shape to the Leithiinae genera *Dryomys* and *Eliomys* (Figure 2). This is presumably the result of the relative broad zygomatic arches and long molar rows in these species. Furthermore, *G. nagtglasii* (*Aethoglis*) is the largest *Graphiurus* species, similar in size with *Eliomys* specimens. The desert dormouse *Selevinia betpakdalaensis* best resembles dormice within Graphiurinae, whereas, phylogenetically, it is placed within the Leithiinae. This is the result of the extremely reduced molars within this specimen, corresponding with the relatively short molar rows seen in *Graphiurus*. The *Selevinia* skull also suggests morphological similarities with *Dryomys* (PC2) and *Myomimus* (PC4), in line with previous studies of this specimen (Hennekam et al., 2020a). In both datasets, the hazel dormouse *Muscardinus avellanarius* is very different compared to other species. This is especially clear when examining shape variation within the mandible. This small arboreal dormouse has highly modified molars, displaying pronounced lateral ridges specialised for eating the tough food it was named after, hazelnuts. Other than this unusual species and the *Graphiurus* subgenera *Claviglis* and *Aethoglis*, the first principal component for skulls and the first two principal components for mandibles distinguish between the three dormouse subfamilies. However, the first principal components do not represent changes in size, even though size variation within the family is substantial. This shows that the allometric effect on shape is relatively small compared to shape variation associated with phylogeny and, potentially, habitat.

### Allometric patterns

Size is a distinguishing feature of dormouse genera (Storch, 1978), and the extent of size variation differs between species (Figures 1 and 4). Analyses indicated the interaction effect of size and genus was significant, and had a smaller residual sum of squares than the analyses without the interaction. However, as the differences between the models was not significant, the common allometric trajectory can be used to evaluate the allometric signals within the datasets (Table S2). This is probably the result of non-significant correlations between shape and size within certain genera. Significant correlations between size and shape are more common in larger dormouse species (*Glis* and *Eliomys*; Table 3). Smaller to medium sized dormice show significant correlations between shape and locality and no significant correlation between shape and size, although this could partly be explained by the relatively small sample size. Shape variation within some genera represents morphological variations between populations. This appears to be the case for *Dryomys* and *Muscardinus*. This could not be testes for other small dormice analysed in this study, as they dataset did not include spatially separated populations for these species (Table 3).

For both datasets, the covariate ‘location’ explains a substantial amount of the total shape variation. Size shows a significant, although smaller, impact on the morphology within dormice as well. The cranial dataset indicates more significant correlations between size and shape compared to the mandibular dataset (Figure 5), with size explaining a larger amount of the total shape variation in the cranium (Table 3). This is also evident in the medium-sized genus *Dryomys*, which shows significant correlations of both size and location with its cranial morphology, whereas location was the only significant variable for the mandibular shape.

The common allometric component for the cranial dataset shows a clear distinction in size between the large dormouse genera (*Glis* and *Eliomys*) and all other genera, with the exception of the peculiar *Graphiurus nagtglasii* (Figure 5). The warps along the allometric trajectory show an elongation of the rostrum, flattening of the cranial vault and relative reduction of the auditory bullae and foramen magnum corresponding with an increase in size. However, the allometric signal in the mandibular dataset is less pronounced. The relatively small genus *Myomimus* actually resembles larger genera (Figure 5). *Muscardinus* appears to be morphologically very different from other small dormice, only resembling the desert dormouse *Selevinia*. Increasing the size of mandibles is associated with a relative shortening of the condylar process and of the diastema. This shape change is very similar to that observed along the third principal component, having the small *Muscardinus* and large *Glis* on either end of the axis (Figure 3). These shape changes are seen to a much lesser extent in the relatively large genus *Eliomys*. This indicates that the allometric trajectory is not just depicting morphological variation associated with size, but could represent shape variations related to genera occupying various ecological niches. The mandibular allometric trajectory is not nearly as well defined compared to the cranial dataset, even though mandibular size is quite variable within dormice.

It appears that although some cranial structures within dormice can be explained by allometry, a variety of features appear to be species-specific. In particular, small dormice vary significantly in morphology, from a very reduced molar row in *Selevinia*, to extremely large molars in *Muscardinus. Selevinia* is also characterised by exceptionally inflated auditory bullae, whereas these are heavily reduced in *Glirulus*. Larger dormouse genera, like *Glis* and *Eliomys* are morphologically very different from each other. However, shape changes within these genera as size increases appear to be relatively similar, indicating similar allometric trajectories. Smaller individuals are characterised by a shortened rostrum and a relatively bulbous cranial vault. Larger specimens show a clear flattening of the cranial vault and a relatively small auditory bulla and foramen magnum. It seems that mandibular morphology in Gliridae is strongly influenced by parameters other than size, including diet, climate and habitat.

### Associations between cranio-mandibular morphology and ecology

#### Size and locomotion

The cranial shape of the three dormouse genera analysed is dependent on at least two variables: size and locality. Size explains only a fraction of the total shape variation (Table 3). However, the shape variation occurring with size variation within groups, especially within species, is consistent. Larger specimens are generally characterized by a flattening of the cranial vault, and presumably a relatively small endocranial volume. This is a pattern often seen in mammalian lineages and associated with the negative allometric scaling of the brain with respect to increased body size (Craniofacial evolutionary allometry “CREA”; Cardini et al., 2015). Furthermore, a more caudal orientation of the foramen magnum is observed in larger specimens, whereas smaller individuals have small robust rostra, an enlarged and rounded cranial vault and a more ventrally oriented foramen magnum. The enlarged cranial vault suggests a relatively larger brain volume, an adaptation generally seen in arboreal mammals (Rensch, 1959; Eisenberg, 1981). The ventral position of the foramen magnum in lagomorphs is believed to be an adaptation to the sitting behavior (DuBrul, 1950), whereas this morphology in hominids is linked with increased use of bipedal locomotion (Russo & Kirk, 2013). However, foramen magnum orientation in non-primates, including rodents, is not always correlated with bipedalism (Russo & Kirk, 2013; Ruth et al. 2016). Satoh & Iwaku (2008) proposed ventral positioning of the foramen in *Apodemus* spp. to be an adaptation to a semi-arboreal niche, in which head posture increases the field of vision. The majority of small-sized dormouse species are considered primarily arboreal, which corresponds with the morphological features seen in smaller specimens. A more caudally orientated foramen magnum is seen in populations of *E. quercinus* from montane woodlands, a region in which this animal is known to be more terrestrial (Holden-Musser et al., 2016).

#### Geographical divergence and habitat specific morphologies

Location explains a large portion of the total shape variation within dormice. However, this variation could be the result of shape variation between species. Then again, speciation within dormice, unsurprisingly, appears to be closely linked with geographical separation of populations and variation in occupied niches. It is therefore of interest to understand the shape variation linked with locality, as it might be indicative for modes of life. Within the family, the shape of the zygomatic arch appears highly variable within species, but only appears to separate species and populations for the genus *Eliomys*, and to some extent in *Graphiurus*. The same applies for the position of the incisive foramina. A morphological feature more clearly linked with ecology is the inflated auditory bullae in specimens associated with dry desert-like environments, as described by Lay (1972) and especially visible within the desert dormouse *Selevinia betpakdalaenis* and certain *Graphiurus* species. Interestingly, the auditory bullae appear not to be inflated in the enlarged *Glis glis* occupying a desert-like habitat (Figure 6), which might indicate that the relative increase in auditory bullae as an adaptation to a desert-like environment is not or less applicable to larger species.

A recurring pattern of cranial vault flattening and a caudally orientated foramen magnum is identified in large terrestrial dormice. The flattening of the skull could be the result of enlarged masticatory muscles in bigger specimens (Penrose et al., 2016) or CREA (Cardini et al., 2015). More agile dormice (arboreal) are relatively small and have a bulbous cranial vault and shortened rostrum. The foramen magnum in these species is orientated ventrally and is relatively large. Similar features are seen in the largest *Graphiurus* species, *G. nagtglasii* (Figure 9), which is documented to be predominantly arboreal and rarely ventures on the ground (Holden-Musser et al., 2016). The smaller, but very terrestrial dormouse, *G. rupicola*, shows clear flattening of the skull and more caudally orientated foramen magnum. The cranial morphology of these large African dormice suggest that the position and relative size of the foramen magnum are indicators of locomotion, and not an artifact of differences in size.

**Figure 9:**
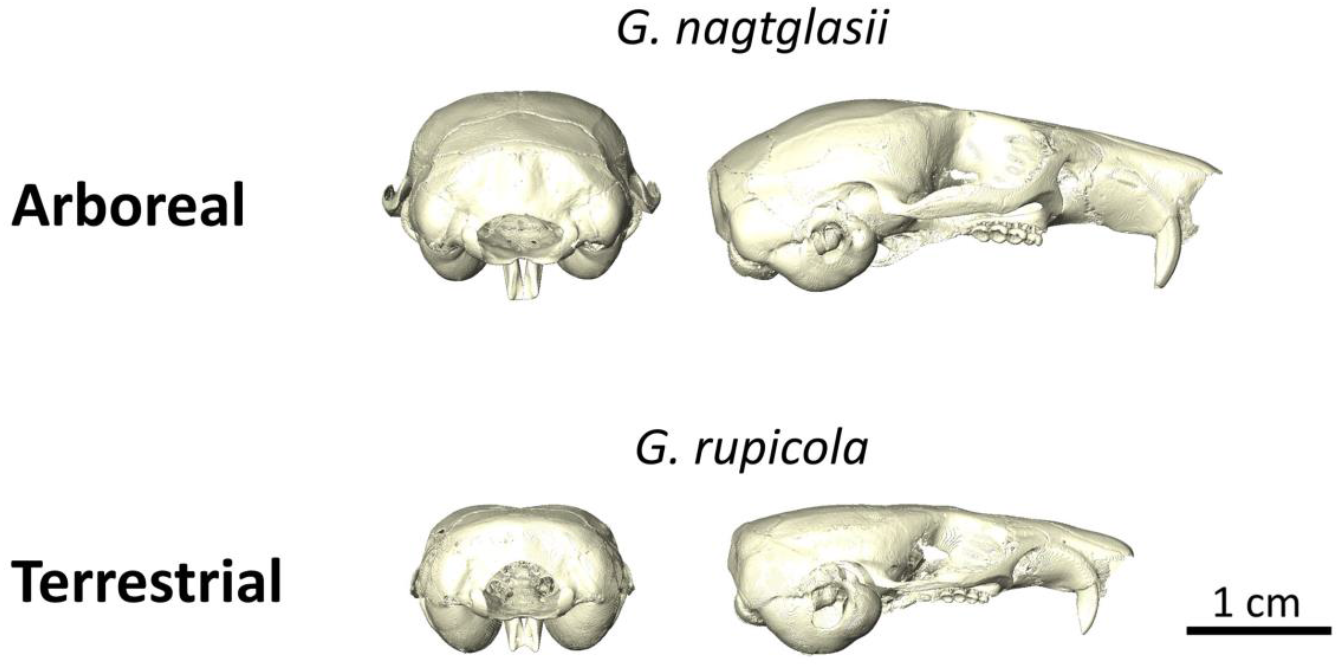
Posterior and lateral view of the arboreal *Graphiurus nagtglasii* and the terrestrial Graphiurus rupicola.

#### Diet related morphologies

Alongside morphologies related to locomotion, some cranial and mandibular shape variations are associated with preferred dietary resources. Landry (1970) stated the ancestral state for Rodentia was omnivory, rather than herbivory, based on the mandible. All species within Gliridae appear to be omnivorous to some degree, although some are clearly more faunivorous or herbivorous than others (Potapova & Rossolimo, 2008; Holden-Musser et al., 2016). The reduced molar rows in *Selevinia* and *Graphiurus ocularis* are thought to be related to a highly insectivorous diet (Webb & Skinner, 1995; Holden-Musser et al., 2016; Hennekam et al., 2020a). *Glis, Glirulus, Muscardinus* and the thick-tailed African dormouse *Graphiurus crassicaudatus* are considered to be the more herbivorous dormice. However, *Glirulus* is known to prefer insects in captivity, suggesting it is more an opportunistic omnivore than a herbivorous dormouse. All other species are simply classified as omnivorous, even though clear variation in diet between species is present (Holden-Musser et al., 2016). The rupicolous dormouse *Graphiurus rupicola* is regarded to be more insectivorous than the arboreal *G. nagtglasii*. Species like *G. murinus* and *E. quercinus* are known to have a variable diet, related to their location (Kahmann & Lau, 1972; Holden-Musser et al., 2016).

More herbivorous dormice are characterised by a relatively robust mandible, with *Muscardinus* displaying an highly unusual mandibular shape for a dormouse (Figures 3 and 5), including a very distinctive molar row. Herbivorous rodents are characterised by a more robust zygomatic arch and a more massive skull in order to withstand the stresses imposed by mastication (Samuels, 2009). In dormice, the zygomatic arch is more robust in herbivorous species like *Glis glis* and *Graphiurus crassicaudatus*, and orientated more horizontally than in most other species. This results in the arch being in line with the occlusal surface of the molar row within these species. Furthermore, pronounced temporal crests are seen in various herbivorous dormice, whereas these are absent in specimens with a more faunivorous diet.

#### Functional implications of morphological variations

Specific morphological features within dormice can be associated with dietary preferences. For example, the angular process of the mandible is positioned more anteriorly in the relatively insectivorous dormouse genera *Graphiurus* and *Selevinia*. The function of this process is associated with the attachment of masseteric muscles on the mandible. The anterior repositioning therefore suggests a reduction and shortening of the deep masseter muscles, which will have implication for the mechanical advantage of the chewing mechanism. A similar positioning of the angular process is seen in omnivorous genera with a preference to insects, including *Glirulus* and *Eliomys*. This feature seems to correspond with a lateral flaring of the coronoid process and a more angled zygomatic arch, resulting a relatively lower glenoid fossa. More insectivorous dormice, like *G. ocularis* and *G. rupicola*, show very thin zygomatic arches. The variation in location of the angular process and the orientation and thickness of the zygomatic arch is presumably connected with the masseter muscle architecture. The more anterior position of the angular process assumes a relative shortening of this muscle, as well as a reduced attachment area on the mandible. The angled zygomatic arch also influences the way this muscle attaches to the skull and potentially the direction of the muscle fibers. A more horizontally orientated zygomatic arch, an elongated angular process, and a less inverted mandibular angular process all result in an enlarged area of muscle attachment for the masseter muscles. The enlarged masseter muscles result in an increase in bite force (Maynard Smith & Savage, 1959). The zygomatic arch and the angular process are more in line with molar rows, which potentially affects the biomechanical efficiency during chewing. These features are appear to be present in more herbivorous dormice.

The lowering of the glenoid fossa and the lateral flaring of the coronoid process seen in relatively insectivorous dormice may have implications on the maximum gape of these animals. As the auditory bullae in dormice are considered inflated with respect to other rodents, these mandibular morphologies might be needed in smaller dormice in order to increase potential gape (Nikolai & Bramble, 1983). Specimens lacking these morphological features, including *Glis*, might therefore not be dependent on an increased potential gape, or are not as constrained by the inflated auditory bullae compared to smaller dormice.

#### Dental morphology

Dentition in dormice is very variable and seems to correlate well with dietary preferences (Wahlert et al., 1993). Small concave and simplistic molars are associated with the more insectivorous species, whereas large robust molars, often including pronounced lateral ridges, are seen in dormice with a more herbivorous diet. The molar row within most *Graphiurus* specimens is relatively small with respect to other genera, with the exception of *Selevinia*, in which this feature is extremely reduced. The two genera are phylogenetically distant from each other and reside in two different parts of the world. *Graphiurus* species occupy a wide range of ecological habitats in Sub-Saharan Africa, whereas the monospecific *Selevinia* is only known from the desert plains in Kazakhstan. However, both genera are considered to be relatively insectivorous (Holden-Musser et al., 2016). Molars in both genera are generally concave, lacking the complex architecture of ridges seen in many other dormouse species (Wahlert et al., 1993; Hennekam et al., 2020a).

The degree of molar development appears to be connected with dietary preference within the highly variable *Graphiurus* genus. The genus is relatively speciose and displays differences in molar characteristics between some populations. Despite being considered omnivorous in general, the dietary preferences within *Graphiurus* span the herbivorous *G. crassicaudatus* to the insectivorous *G. ocularis* and *G. rupicola* species. More herbivorous species show clear adaptation to their diets: *G. nagtglasii* appears to be more herbivorous than most other *Graphiurus* species and displays more highly developed molar ridges (Figure 10). These ridges are even more pronounced in the predominantly herbivorous *G. crassicaudatus*, but lacking in the more omnivorous and insectivorous species (e.g. *G. murinus, G. rupicola*). The patterns seen in *Graphiurus* resemble morphological adaptations in other genera, with the extremely reduced molars in the insectivorous *G. ocularis* and the desert dormouse *Selevinia*, and more developed molar structures in herbivorous specimens such as *G. crassicaudatus* and genera like *Glis* and *Muscardinus*.

**Figure 10:**
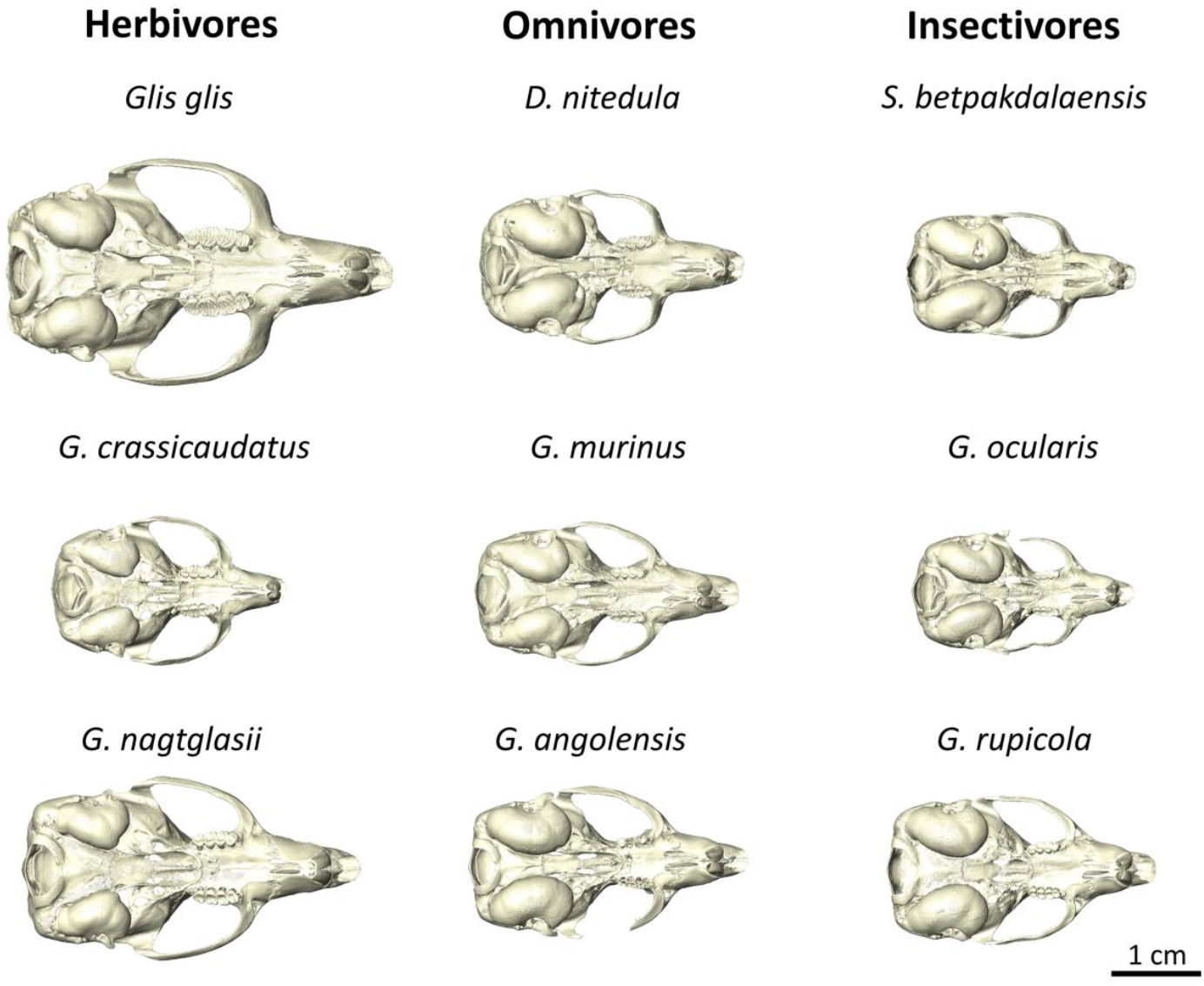
*Ventral orientation of dormice allocated to specific dietary ecologies. Genus abbreviations: D =* Dryomys; *S =* Selevinia; *G =* Graphiurus.

### Conclusion

The variation in cranial and mandibular morphology of eight out of nine extant dormouse genera was evaluated. Dormice appear to vary in shape and size significantly between genera, and to a lesser extent between and within species. Larger dormice are more variable in size than smaller dormouse genera, with the exception of the speciose genus *Graphiurus*. The common allometric component identifies a short rostrum, a bulbous cranial vault and a large ventrally orientated foramen magnum as characteristic for smaller dormice, whereas flattening of the cranial vault and a more caudally positioned foramen are seen in larger specimens. Mandibular morphology is less driven by allometry, but does show relatively enlarged angular processes in larger dormice. Specific morphologies seem to be associated with specific locomotor and dietary ecologies. The position of the foramen magnum can be used as an indication for a more arboreal or terrestrial lifestyle, whereas the orientation and thickness of the zygomatic arch, combined with the relative size and morphology of the molar row are related to diet. This study connects the large variety of habitats and behaviours in dormice with morphological features, and clearly highlights the strong relationship between form and function in the mammalian skull.

## Acknowledgements

The author would like to thank Philip Cox, for his comments and guidance on the study. Nathan Jeffery and Roger Benson for providing operable scan data. Tom Davies at the scanning facility in Bristol. Victoria Herridge for PhD supervision and advice. Irina Ruf (SMF), Roberto Portela-Miguez (NHMUK), and Violaine Nicolas (MNHM) for providing the dormouse specimens.

## Data Availability Statement

The 3D scans and reconstructions used in this study were uploaded onto the MorphoSource online repository: morphosource.org/projects/00000C941

## Supplementary Information

**Figure S1:**
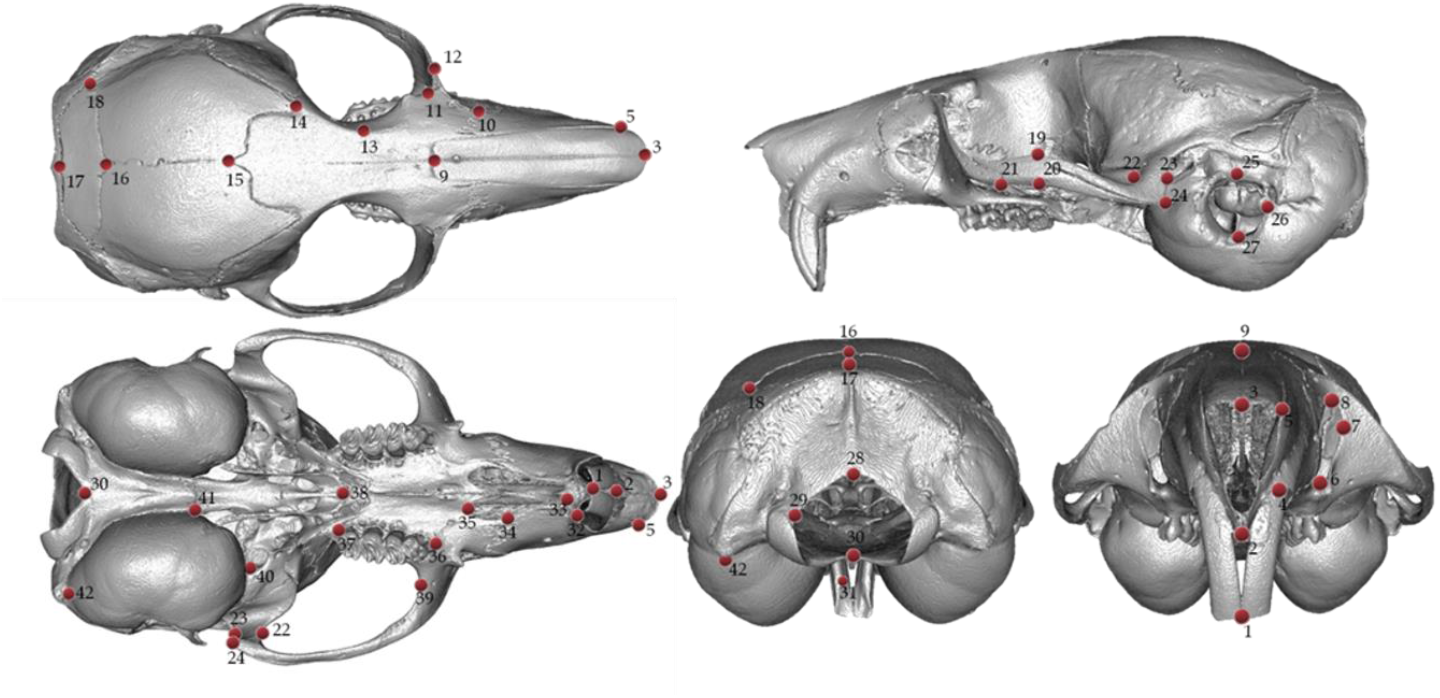
Cranial landmark configuration consisting of 42 anatomical landmarks, visualised in dorsal, lateral, ventral, posterior and anterior view.

**Figure S2:**
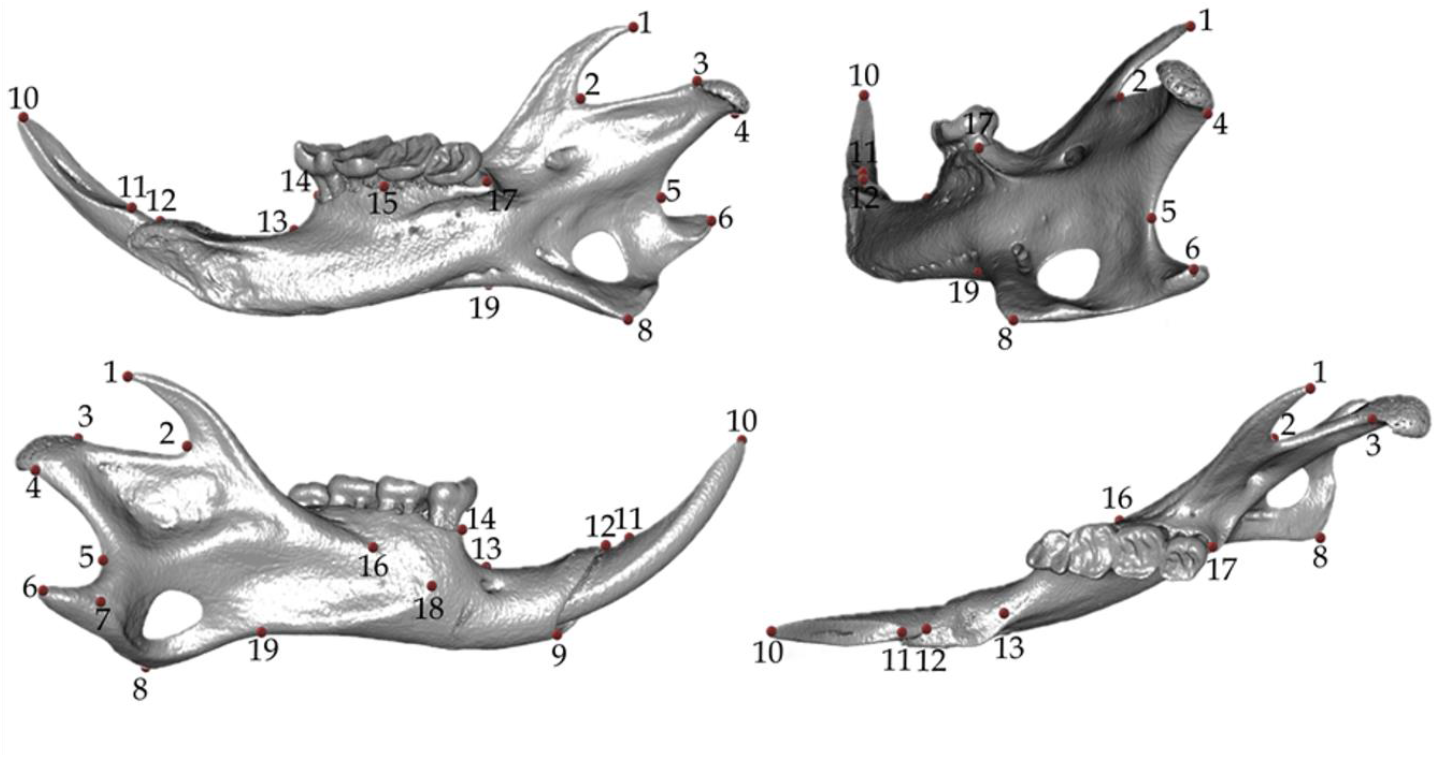
Mandibular landmark configuration consisting of 19 anatomical landmarks, visualised in medial, posterior, lateral and dorsal view.

**Figure S3:**
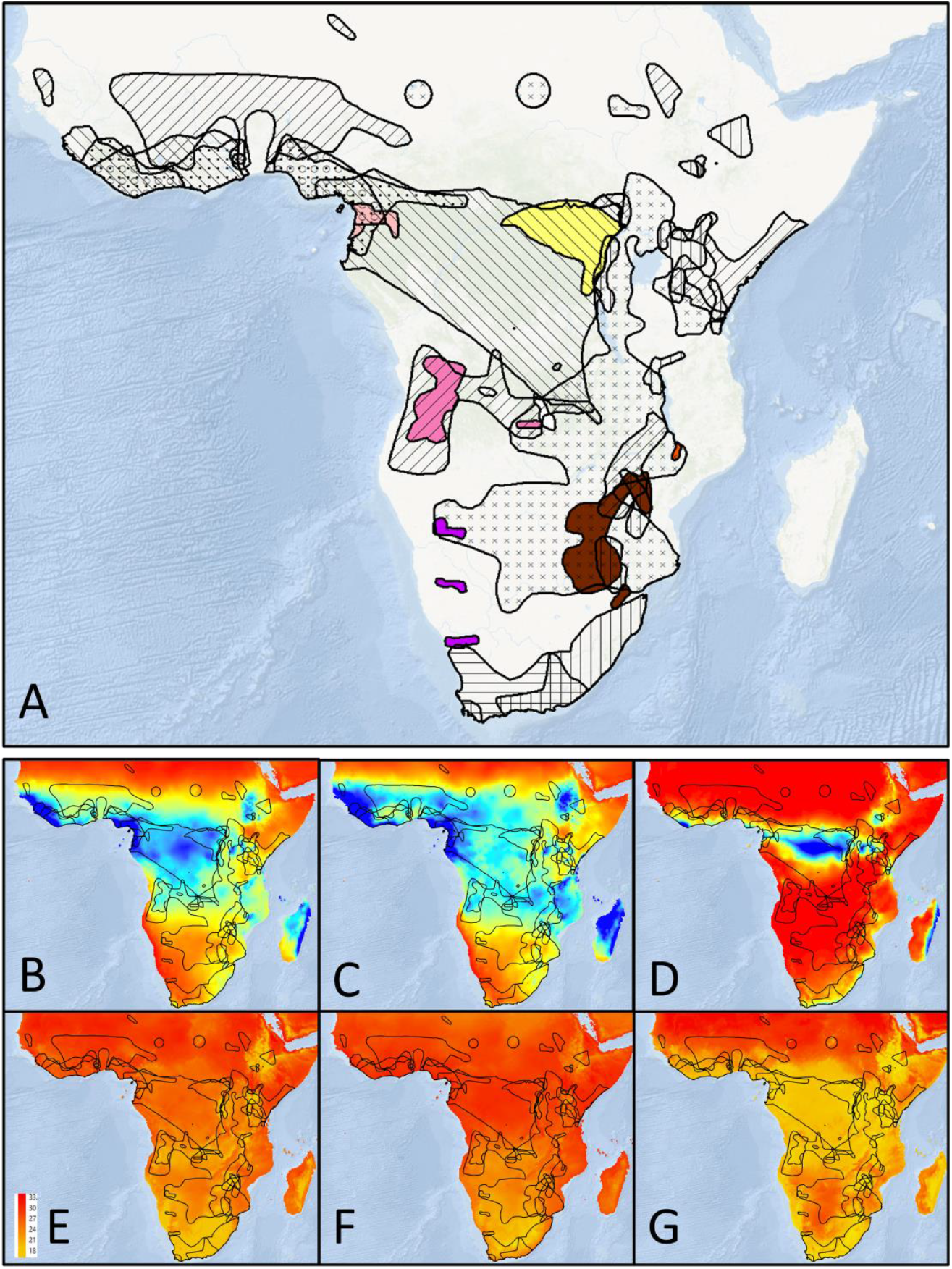
Dispersal of the genus *Graphiurus* in sub-Saharan Africa (A) according to IUCN. B – G include the outline of the dispersal areas and display climatic data derived from WorldClim v2. B-D project a heat map indicating Mean Annual Precipitation (B), precipitation in the wettest month (C) and in the driest month (D). E-G show display temperature differences: mean annual temperature (E), mean temperature of the warmest month (F) and of the coldest month (G).

**Figure S4:**
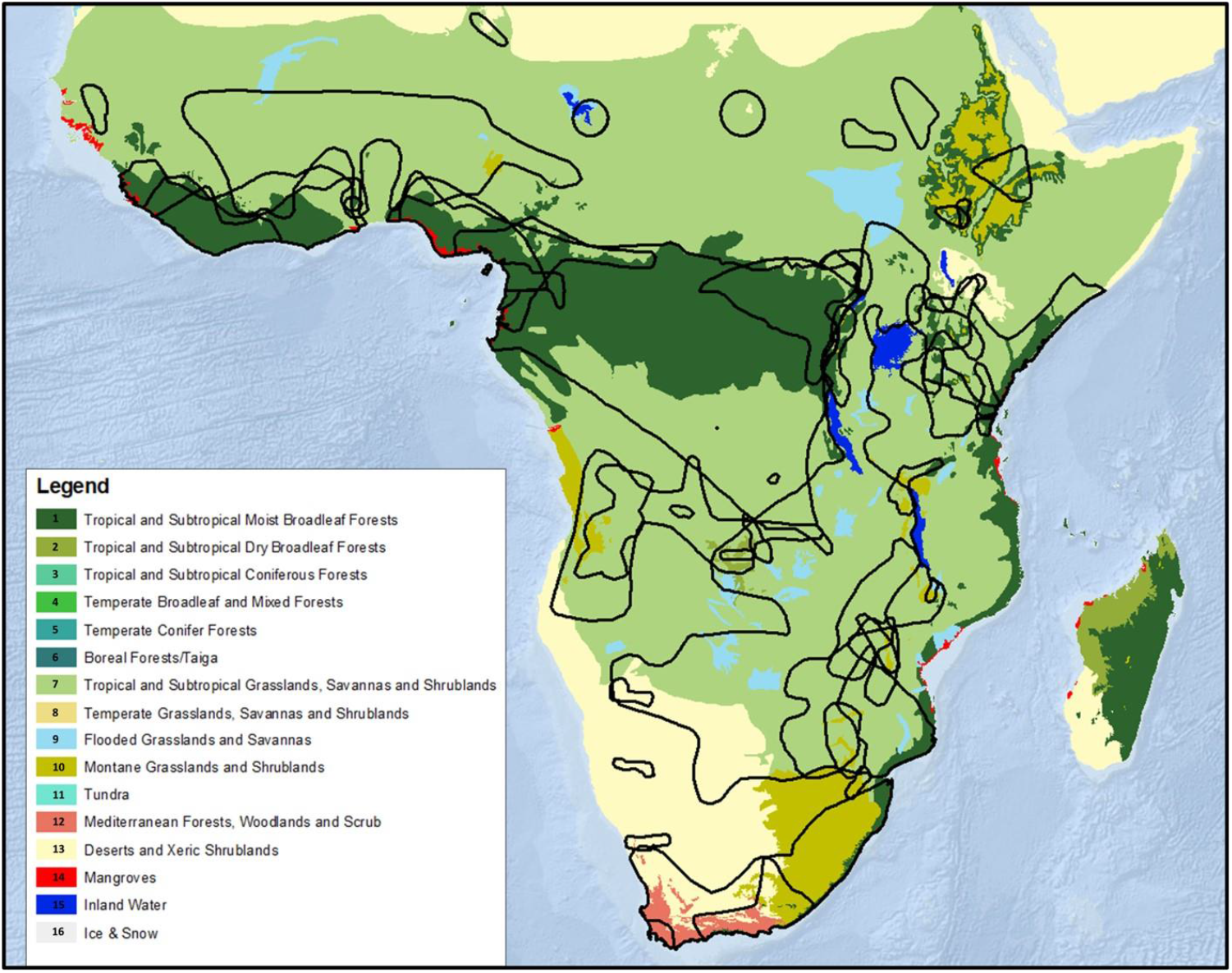
Ecoregions in sub-Saharan Africa according to the WWF, including the outlines of the dispersal areas for *Graphiurus*.

**Table S1:**
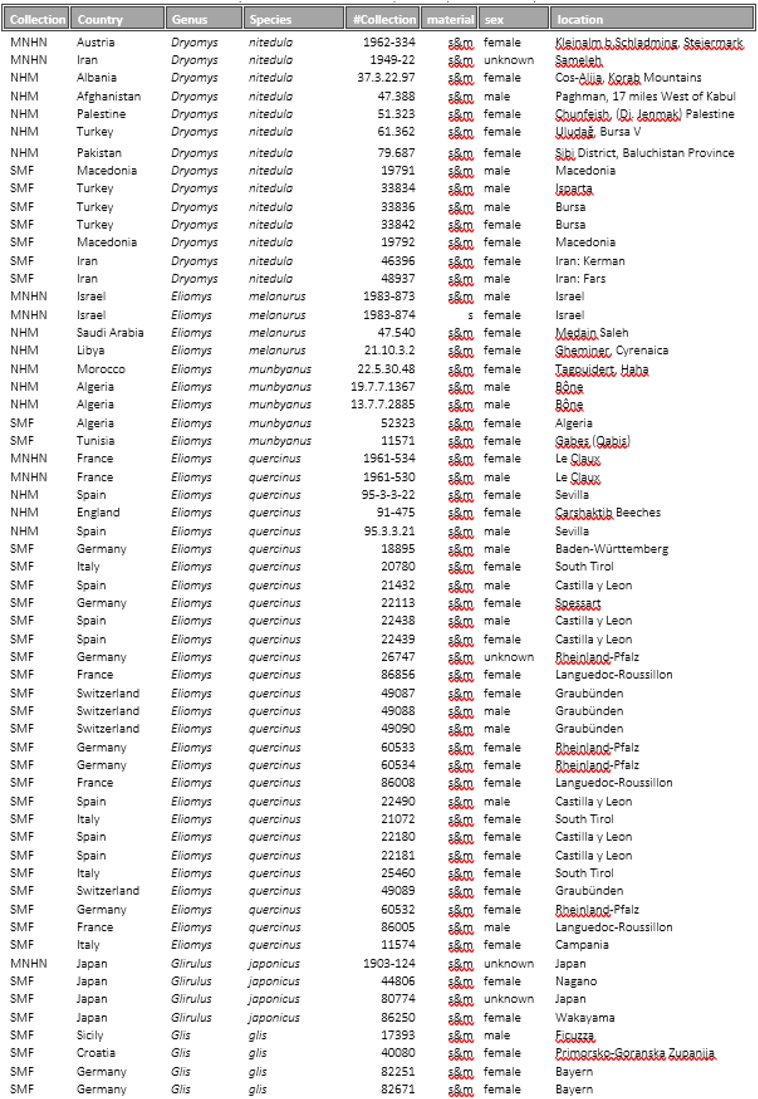
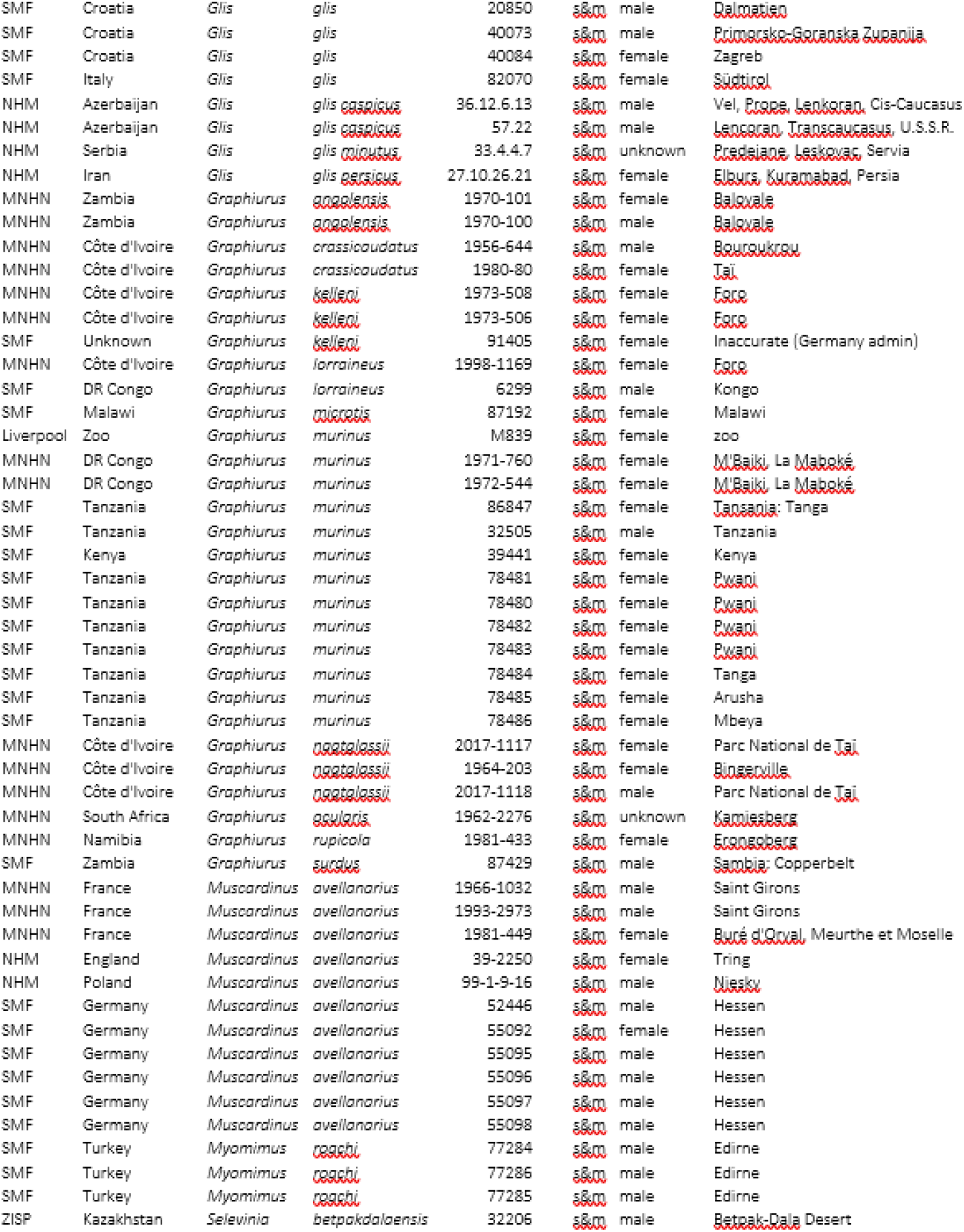
List of all dormouse specimens used in shape analyses for Chapter 3.

**Table S2:**
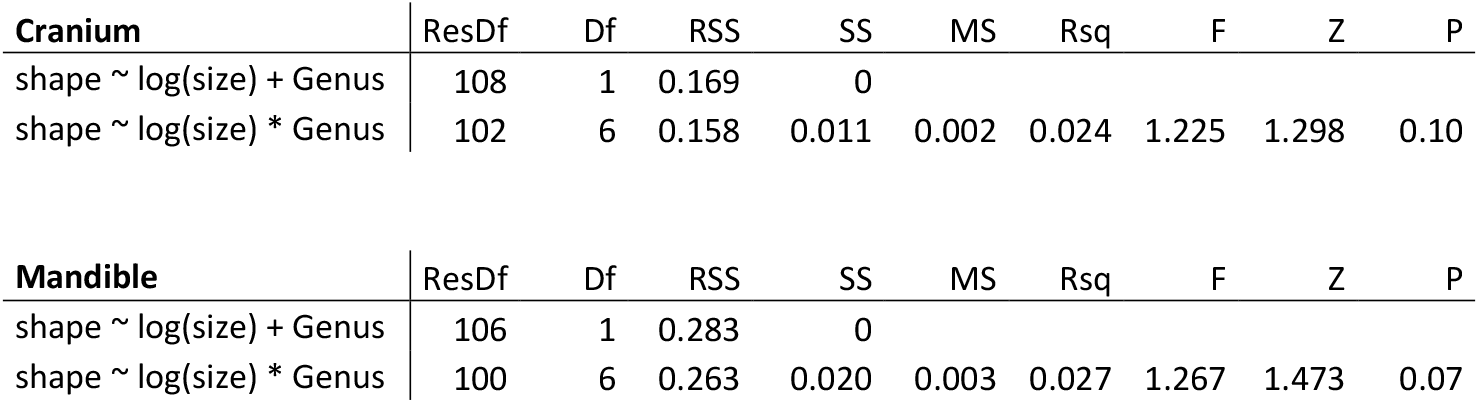
Testing for unique or common allometries within the skull (top) and mandibular dataset (bottom).

**Table S3:**
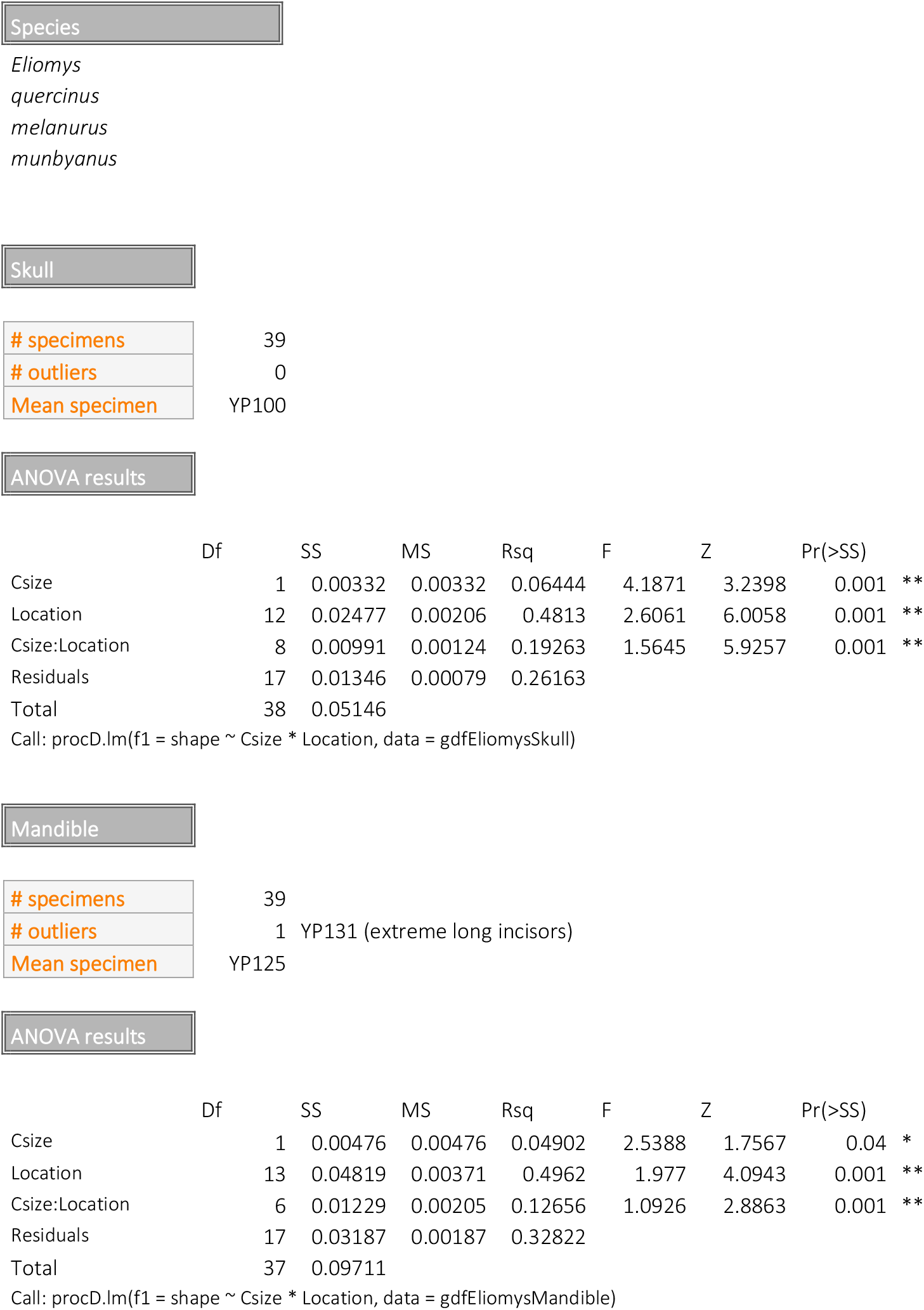
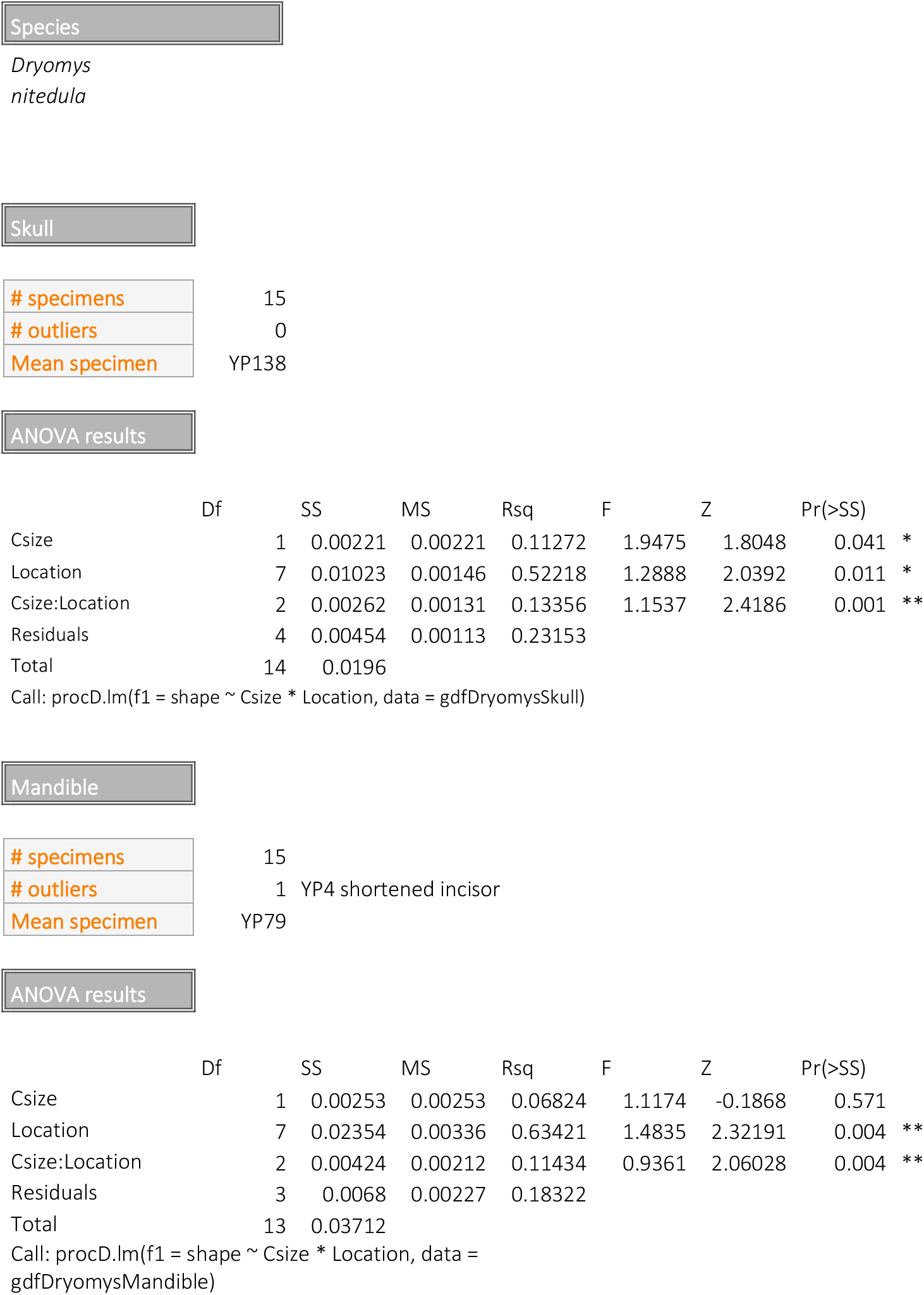
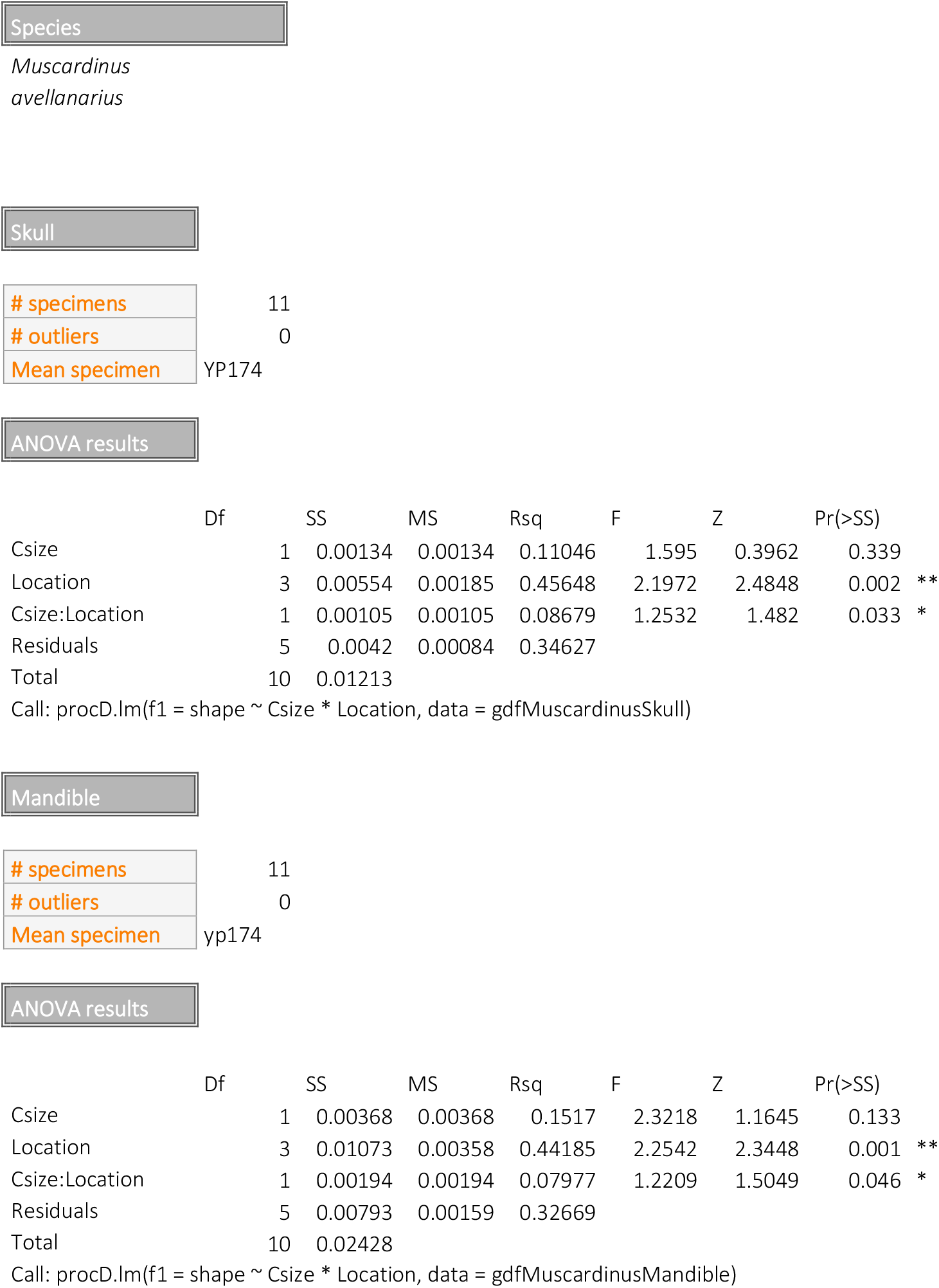
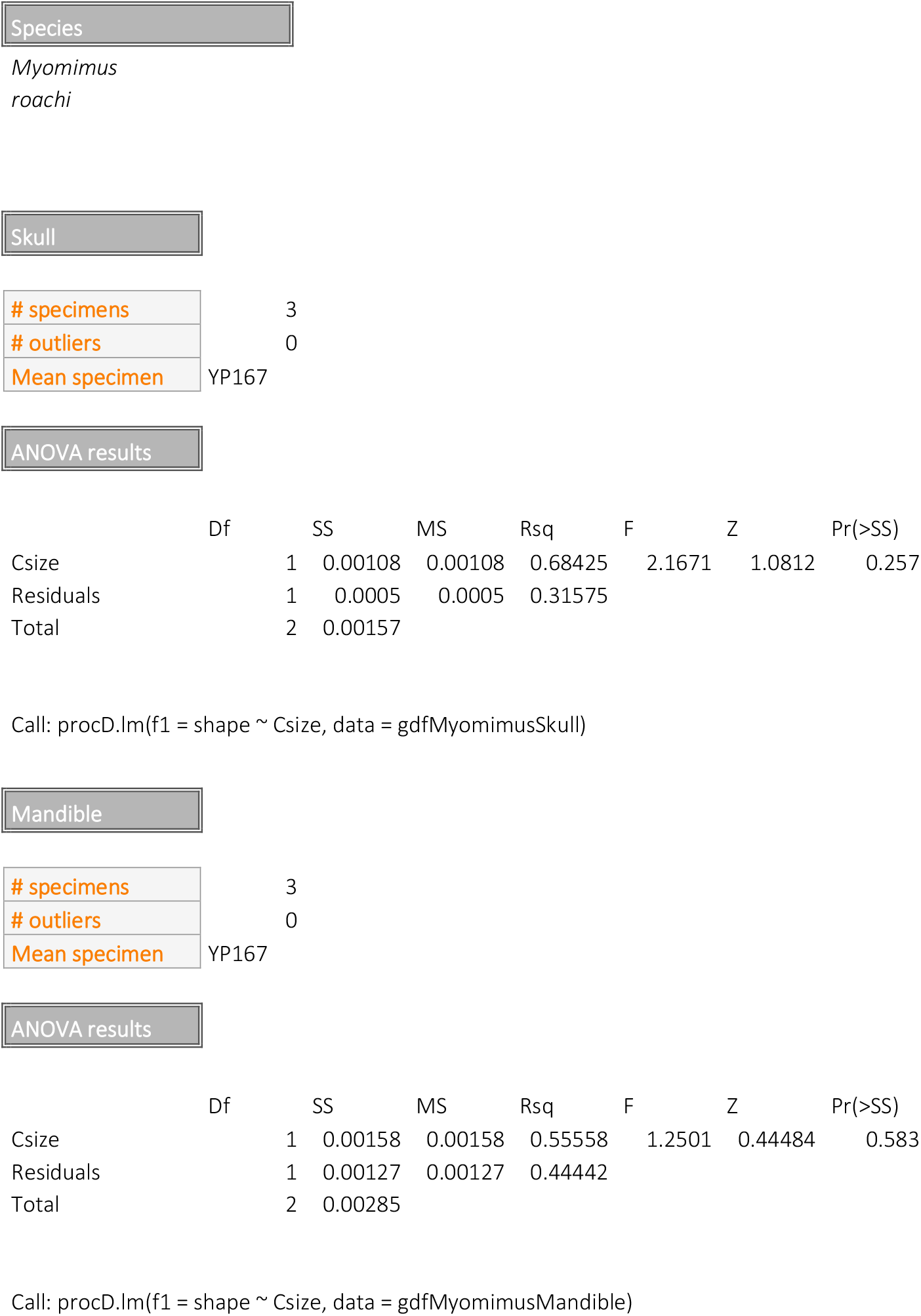
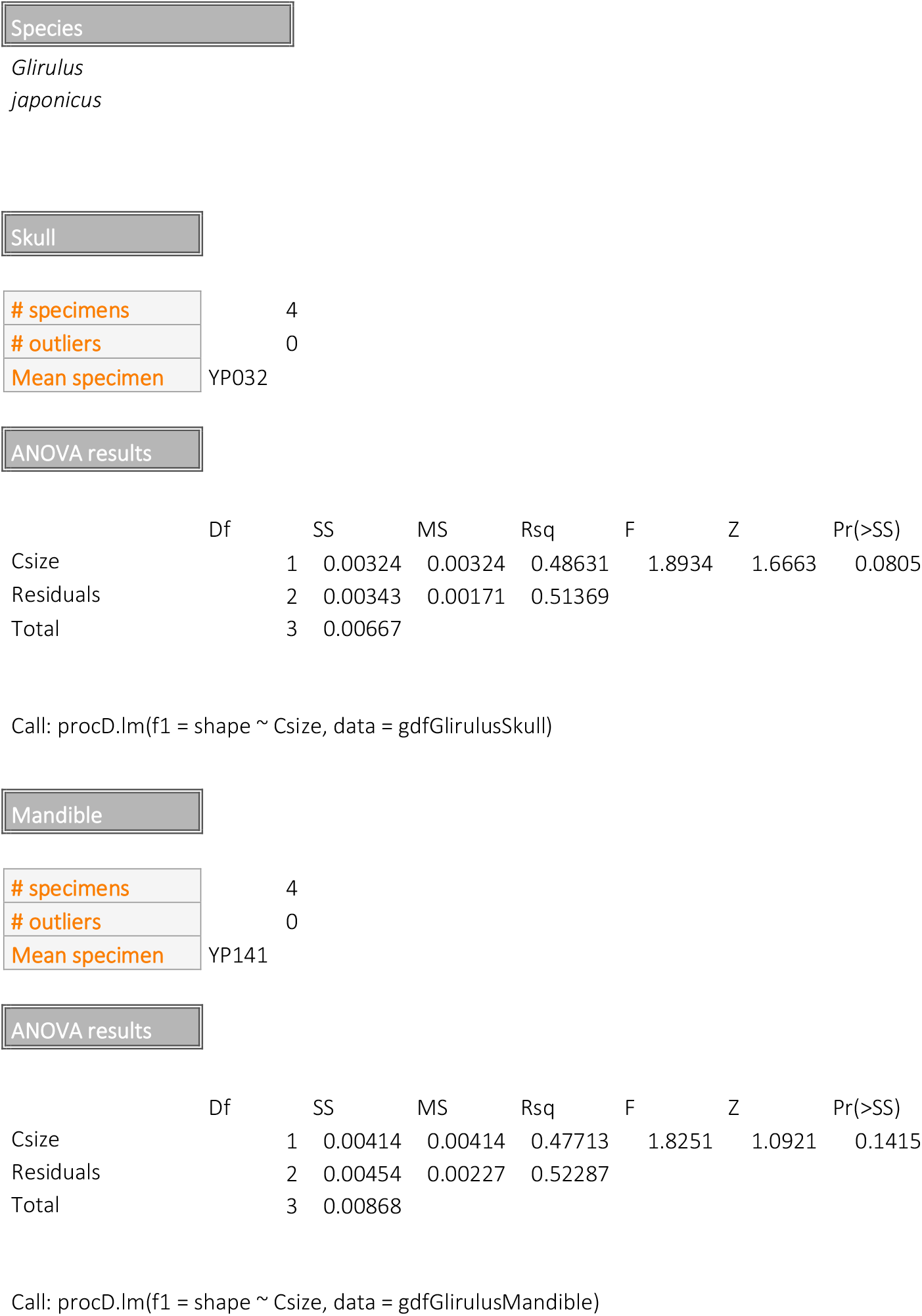
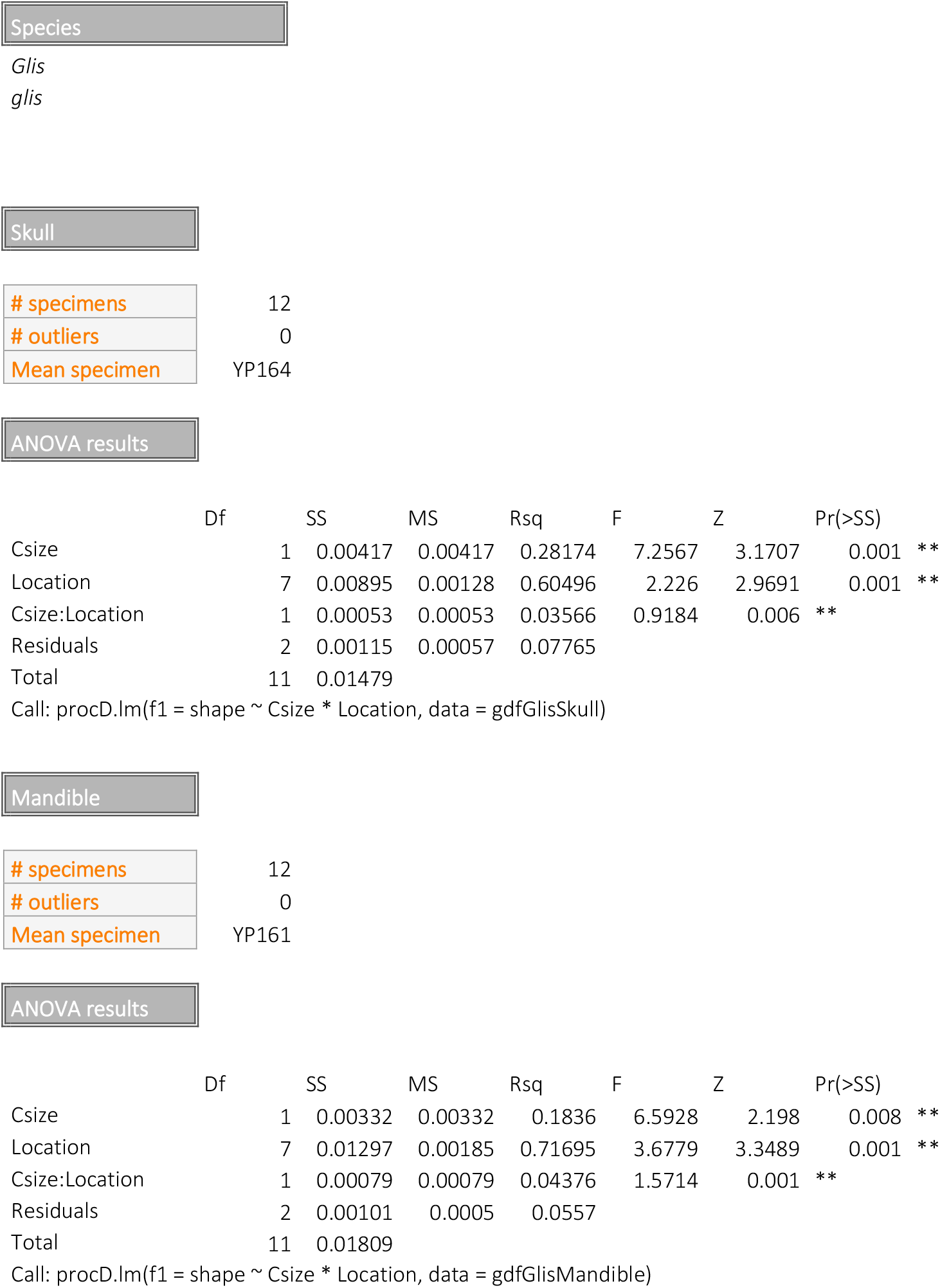
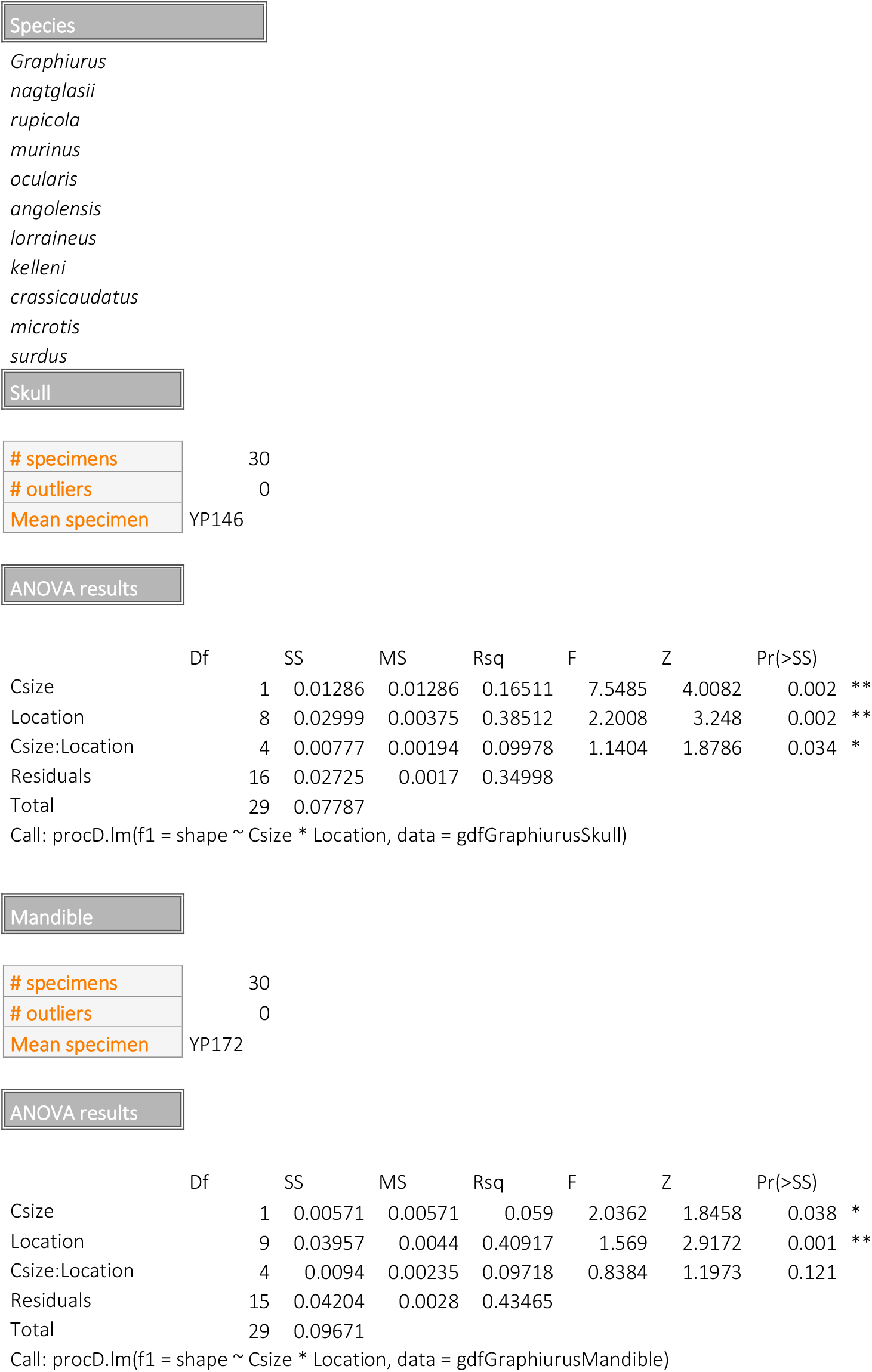
Procrustes ANOVAs per genus

|  |  |
| --- | --- |
| # specimens | 39 |
| # outliers | 0 |
| Mean specimen | YP100 |

ANOVA results
|  | Df | SS | MS | Rsqr | F | Z | Pr(>SS) |  |
| --- | --- | --- | --- | --- | --- | --- | --- | --- |
| Csize | 1 | 0.00332 | 0.00332 | 0.06444 | 4.1871 | 3.2398 | 0.001 | ** |
| Location | 12 | 0.02477 | 0.00206 | 0.4813 | 2.6061 | 6.0058 | 0.001 | ** |
| Csize:Location | 8 | 0.00991 | 0.00124 | 0.19263 | 1.5645 | 5.9257 | 0.001 | ** |
| Residuals | 17 | 0.01346 | 0.00079 | 0.26163 |  |  |  |  |
| Total | 38 | 0.05146 |  |  |  |  |  |  |
| # specimens | 39 |
| # outliers | 1 YP131 (extreme long incisors) |
| Mean specimen | YP125 |

ANOVA results
|  | Df | SS | MS | Rsqr | F | Z | Pr(>SS) |  |
| --- | --- | --- | --- | --- | --- | --- | --- | --- |
| Csize | 1 | 0.00476 | 0.00476 | 0.04902 | 2.5388 | 1.7567 | 0.04 | * |
| Location | 13 | 0.04819 | 0.00371 | 0.4962 | 1.977 | 4.0943 | 0.001 | ** |
| Csize:Location | 6 | 0.01229 | 0.00205 | 0.12656 | 1.0926 | 2.8863 | 0.001 | ** |
| Residuals | 17 | 0.03187 | 0.00187 | 0.32822 |  |  |  |  |
| Total | 37 | 0.09711 |  |  |  |  |  |  |
| Skull |  |
| # specimens | 15 |
| # outliers | 0 |
| Mean specimen | YP138 |

ANOVA results
|  | Df | SS | MS | Rsqr | F | Z | Pr(>SS) |
| --- | --- | --- | --- | --- | --- | --- | --- |
| Csize | 1 | 0.00221 | 0.00221 | 0.11272 | 1.9475 | 1.8048 | 0.041 * |
| Location | 7 | 0.01023 | 0.00146 | 0.52218 | 1.2888 | 2.0392 | 0.011 * |
| Csize:Location | 2 | 0.00262 | 0.00131 | 0.13356 | 1.1537 | 2.4186 | 0.001 ** |
| Residuals | 4 | 0.00454 | 0.00113 | 0.23153 |  |  |  |
| Total | 14 | 0.0196 |  |  |  |  |  |
| Mandible |  |
| # specimens | 15 |
| # outliers | 1 YP4 shortened incisor |
| Mean specimen | YP79 |

ANOVA results
|  | Df | SS | MS | Rsqr | F | Z | Pr(>SS) |
| --- | --- | --- | --- | --- | --- | --- | --- |
| Csize | 1 | 0.00253 | 0.00253 | 0.06824 | 1.1174 | -0.1868 | 0.571 |
| Location | 7 | 0.02354 | 0.00336 | 0.63421 | 1.4835 | 2.32191 | 0.004 ** |
| Csize:Location | 2 | 0.00424 | 0.00212 | 0.11434 | 0.9361 | 2.06028 | 0.004 ** |
| Residuals | 3 | 0.0068 | 0.00227 | 0.18322 |  |  |  |
| Total | 13 | 0.03712 |  |  |  |  |  |
| Skull |  |
| # specimens | 11 |
| # outliers | 0 |
| Mean specimen | YP174 |

ANOVA results
|  | Df | SS | MS | Rsqr | F | Z | Pr(>SS) |
| --- | --- | --- | --- | --- | --- | --- | --- |
| Csize | 1 | 0.00134 | 0.00134 | 0.11046 | 1.595 | 0.3962 | 0.339 |
| Location | 3 | 0.00554 | 0.00185 | 0.45648 | 2.1972 | 2.4848 | 0.002 ** |
| Csize:Location | 1 | 0.00105 | 0.00105 | 0.08679 | 1.2532 | 1.482 | 0.033 * |
| Residuals | 5 | 0.0042 | 0.00084 | 0.34627 |  |  |  |
| Total | 10 | 0.01213 |  |  |  |  |  |
| Mandible |  |
| # specimens | 11 |
| # outliers | 0 |
| Mean specimen | yp174 |

ANOVA results
|  | Df | SS | MS | Rsqr | F | Z | Pr(>SS) |
| --- | --- | --- | --- | --- | --- | --- | --- |
| Csize | 1 | 0.00368 | 0.00368 | 0.1517 | 2.3218 | 1.1645 | 0.133 |
| Location | 3 | 0.01073 | 0.00358 | 0.44185 | 2.2542 | 2.3448 | 0.001 ** |
| Csize:Location | 1 | 0.00194 | 0.00194 | 0.07977 | 1.2209 | 1.5049 | 0.046 * |
| Residuals | 5 | 0.00793 | 0.00159 | 0.32669 |  |  |  |
| Total | 10 | 0.02428 |  |  |  |  |  |
| Skull |  |
| # specimens | 3 |
| # outliers | 0 |
| Mean specimen | YP167 |

ANOVA results
|  | Df | SS | MS | Rsq | F | Z | Pr(>SS) |
| --- | --- | --- | --- | --- | --- | --- | --- |
| Csize | 1 | 0.00108 | 0.00108 | 0.68425 | 2.1671 | 1.0812 | 0.257 |
| Residuals | 1 | 0.0005 | 0.0005 | 0.31575 |  |  |  |
| Total | 2 | 0.00157 |  |  |  |  |  |

ANOVA results
|  | Df | SS | MS | Rsq | F | Z | Pr(>SS) |
| --- | --- | --- | --- | --- | --- | --- | --- |
| Csize | 1 | 0.00158 | 0.00158 | 0.55558 | 1.2501 | 0.44484 | 0.583 |
| Residuals | 1 | 0.00127 | 0.00127 | 0.44442 |  |  |  |
| Total | 2 | 0.00285 |  |  |  |  |  |
| Skull |  |
| # specimens | 4 |
| # outliers | 0 |
| Mean specimen | YP032 |

ANOVA results
|  | Df | SS | MS | Rsq | F | Z | Pr(>SS) |
| --- | --- | --- | --- | --- | --- | --- | --- |
| Csize | 1 | 0.00324 | 0.00324 | 0.48631 | 1.8934 | 1.6663 | 0.0805 |
| Residuals | 2 | 0.00343 | 0.00171 | 0.51369 |  |  |  |
| Total | 3 | 0.00667 |  |  |  |  |  |
| Mandible |  |
| # specimens | 4 |
| # outliers | 0 |
| Mean specimen | YP141 |

ANOVA results
|  | Df | SS | MS | Rsq | F | Z | Pr(>SS) |
| --- | --- | --- | --- | --- | --- | --- | --- |
| Csize | 1 | 0.00414 | 0.00414 | 0.47713 | 1.8251 | 1.0921 | 0.1415 |
| Residuals | 2 | 0.00454 | 0.00227 | 0.52287 |  |  |  |
| Total | 3 | 0.00868 |  |  |  |  |  |
| Skull |  |
| # specimens | 12 |
| # outliers | 0 |
| Mean specimen | YP164 |

ANOVA results
|  | Df | SS | MS | Rsqr | F | Z | Pr(>SS) |
| --- | --- | --- | --- | --- | --- | --- | --- |
| Csize | 1 | 0.00417 | 0.00417 | 0.28174 | 7.2567 | 3.1707 | 0.001 ** |
| Location | 7 | 0.00895 | 0.00128 | 0.60496 | 2.226 | 2.9691 | 0.001 ** |
| Csize:Location | 1 | 0.00053 | 0.00053 | 0.03566 | 0.9184 | 0.006 | ** |
| Residuals | 2 | 0.00115 | 0.00057 | 0.07765 |  |  |  |
| Total | 11 | 0.01479 |  |  |  |  |  |
| Mandible |  |
| # specimens | 12 |
| # outliers | 0 |
| Mean specimen | YP161 |

ANOVA results
|  | Df | SS | MS | Rsqr | F | Z | Pr(>SS) |
| --- | --- | --- | --- | --- | --- | --- | --- |
| Csize | 1 | 0.00332 | 0.00332 | 0.1836 | 6.5928 | 2.198 | 0.008 ** |
| Location | 7 | 0.01297 | 0.00185 | 0.71695 | 3.6779 | 3.3489 | 0.001 ** |
| Csize:Location | 1 | 0.00079 | 0.00079 | 0.04376 | 1.5714 | 0.001 | ** |
| Residuals | 2 | 0.00101 | 0.0005 | 0.0557 |  |  |  |
| Total | 11 | 0.01809 |  |  |  |  |  |
| Skull |  |
| # specimens | 30 |
| # outliers | 0 |
| Mean specimen | YP146 |

ANOVA results
|  | Df | SS | MS | Rsqr | F | Z | Pr(>SS) |  |
| --- | --- | --- | --- | --- | --- | --- | --- | --- |
| Csize | 1 | 0.01286 | 0.01286 | 0.16511 | 7.5485 | 4.0082 | 0.002 | ** |
| Location | 8 | 0.02999 | 0.00375 | 0.38512 | 2.2008 | 3.248 | 0.002 | ** |
| Csize:Location | 4 | 0.00777 | 0.00194 | 0.09978 | 1.1404 | 1.8786 | 0.034 | * |
| Residuals | 16 | 0.02725 | 0.0017 | 0.34998 |  |  |  |  |
| Total | 29 | 0.07787 |  |  |  |  |  |  |
| Mandible |  |
| # specimens | 30 |
| # outliers | 0 |
| Mean specimen | YP172 |

ANOVA results
|  | Df | SS | MS | Rsqr | F | Z | Pr(>SS) |  |
| --- | --- | --- | --- | --- | --- | --- | --- | --- |
| Csize | 1 | 0.00571 | 0.00571 | 0.059 | 2.0362 | 1.8458 | 0.038 | * |
| Location | 9 | 0.03957 | 0.0044 | 0.40917 | 1.569 | 2.9172 | 0.001 | ** |
| Csize:Location | 4 | 0.0094 | 0.00235 | 0.09718 | 0.8384 | 1.1973 | 0.121 |  |
| Residuals | 15 | 0.04204 | 0.0028 | 0.43465 |  |  |  |  |
| Total | 29 | 0.09671 |  |  |  |  |  |  |

Call: procD.lm(f1 = shape ~ Csize \* Location, data = gdfGraphiurusMandible)

## Notes

### Competing Interest Statement

The authors have declared no competing interest.

